# Optical Control of Cell-Surface and Endomembrane-Exclusive β-Adrenergic Receptor Signaling

**DOI:** 10.1101/2024.02.14.580335

**Authors:** Waruna Thotamune, Sithurandi Ubeysinghe, Kendra K. Shrestha, Mahmoud Elhusseiny Mostafa, Michael C. Young, Ajith Karunarathne

## Abstract

Beta-adrenergic receptors (βARs) are G protein-coupled receptors (GPCRs) that mediate catecholamine-induced stress responses, such as heart rate increase and bronchodilation. In addition to signals from the cell surface, βARs also broadcast non-canonical signaling activities from the cell interior membranes (endomembranes). Dysregulation of these receptor pathways underlies severe pathological conditions. Excessive βAR stimulation is linked to cardiac hypertrophy, leading to heart failure, while impaired stimulation causes compromised fight or flight stress responses and homeostasis. In addition to plasma membrane βAR, emerging evidence indicates potential pathological implications of deeper endomembrane βARs, such as inducing cardiomyocyte hypertrophy and apoptosis, underlying heart failure. However, the lack of approaches to control their signaling in subcellular compartments exclusively has impeded linking endomembrane βAR signaling with pathology. Informed by the β1AR-catecholamine interactions, we engineered an efficiently photo-labile, protected hydroxy β1AR pro-ligand (OptoIso) to trigger βAR signaling at the cell surface, as well as exclusive endomembrane regions upon blue light stimulation. Not only does OptoIso undergo blue light deprotection in seconds, but it also efficiently enters cells and allows examination of G protein heterotrimer activation exclusively at endomembranes. In addition to its application in the optical interrogation of βARs in unmodified cells, given its ability to control deep organelle βAR signaling, OptoIso will be a valuable experimental tool.

## 1. INTRODUCTION

Approximately one-third to half of all currently available prescription drugs target G protein coupled receptors (GPCRs), indicating their pathophysiological significance.^1^ Beta-adrenergic receptors (βARs) are a subset of GPCRs found in different cells throughout the body.^2^ They are activated by catecholamines such as epinephrine (adrenaline) and norepinephrine (noradrenaline), released by the sympathetic nervous system upon experiencing stress stimuli.^3^ The members of βARs: β1, β2, and β3, show distinct distributions in the body and play numerous physiological roles.^4^ The heart is the primary location of β1ARs and regulate cardiac contractility, heart rate, and conduction velocity.^5^ β2ARs are found in tissues such as bronchial, vascular smooth, and skeletal muscles, as well as cardiac muscles.^6^ They induce smooth muscle relaxation, control vasodilation, and increase glycogenolysis and lipolysis in skeletal muscle.^7^ β3ARs are expressed in adipose tissues, where they regulate lipolysis and thermogenesis.^8^ At the cellular level, upon stimulation by catecholamines all three βARs primarily bind to stimulatory G (Gs) proteins and increase intracellular cyclic adenosine monophosphate (cAMP) levels, activating protein kinase A (PKA).^9^ PKA then phosphorylates troponin I, the L-type calcium channel, and phospholamban, increasing myocellular calcium entry and the release of calcium from the sarcoplasmic reticulum.^10^ The released calcium interacts with the myocardial contractile machinery and produces systole, resulting in a higher contractility.^11^ βAR signaling has distinct post-agonist stimulation regulatory mechanisms such as deactivation of G-proteins after the G protein heterotrimer dissociation and receptor desensitization, to prevent overstimulation. The desensitization begins when the Gβγ subunit binds with the active form of a G-protein-coupled receptor kinase (GRK). GRK2 is the most prominent cardiac GRK, while GRK3 and GRK5 are expressed in low levels but are also capable of phosphorylating the C terminus of βARs.^12^ This phosphorylation induces the arrestin-induced functional uncoupling from the G protein, preventing the overactivation of βARs.^13^ Following desensitization, βARs undergo another important phenomena known as resensitization regulated by various proteins and cellular processes. Once internalized, βARs are recycled back to the cell membrane through the actions of proteins such as Rab proteins, dynamin and phosphatase 2A in the early endosomes, or they may undergo degradation in lysosomes.^14^ During the recycling process, accessory proteins like sorting nexins and the retromer complex play crucial roles in directing the receptors back to the cell surface.^15, 16^ This orchestrated interplay of proteins and cellular events ensures the dynamic regulation of βAR responsiveness, contributing to the maintenance of cellular homeostasis in response to various physiological and pathological conditions such as asthma.^17, 18^

Dysregulation of βARs leads to severe pathologies. For instance, β1ARs are central to heart failure due to their aberrant signaling associated with one or many cellular conditions such as hypertrophy and apoptosis.^12^ At the molecular level, myocardial βAR dysfunction in heart failure is characterized by loss of β1AR density at the plasma membrane and by the desensitization of β1ARs and β2ARs.^19^ Sustained activation of β2ARs is linked to bronchial hyperreactivity found in asthma.^20^ Impaired signaling of both β2 and β3ARs are involved in metabolic disorders such as obesity and diabetes.^21^ Therefore, agonists that activate βARs are drugs that control many physiological responses, such as increasing heart rate and cardiac output, bronchodilational respiratory conditions (including asthma), and chronic obstructive pulmonary disease (COPD). Beta-blockers are commonly known drugs that act as antagonists for βARs. To date, three generations of beta blockers have been released as drugs.^22^ The first generation consists of nonselective β blockers while the second generation contains cardio selective β blockers that are selective β1AR antagonists.^22^ The third generation of these drugs are able to block β1Ars together with extra vasodilation activity either by blocking α1 or by activating β3AR.^22^ These drugs are extensively used to treat various conditions, including hypertension, angina, arrhythmias, and heart failure.^22^ By blocking the activity of β1ARs in the heart, beta-blockers can reduce heart rate and cardiac output, which can be beneficial in these conditions.^22^

Though GPCR-G protein signaling was thought to be restricted to the plasma membrane, emerging evidence shows GPCR activation and signaling in endomembrane locations such as Golgi, endoplasmic reticulum (ER), and the nuclear membrane.^23^ The presence of GPCRs in the nuclear membranes of neurons and cardiomyocytes,^24, 25^ and βARs in their active conformation in Golgi have been demonstrated.^26^ Also, the link between cardiac hypertrophy and β1AR activity in Golgi has been demonstrated.^27^ Furthermore, it is documented that the exclusive activation of endomembrane G proteins regulates cardiomyocyte hypertrophy which causes an abnormal enlargement or thickening in heart muscles.^28^ Interestingly, it has been shown that, β1ARs expressed in endomembranes like Golgi exhibit an involvement in cardiomyocyte hypertrophy and apoptosis while β2ARs expressed in early endosomes closer to the plasma membrane, exhibit cardioprotective activity by alleviating the β1AR effects^29^. Ligand-bound internalized receptors in endocytic vesicles and *de novo* GPCRs near Golgi, activated by cell-permeable ligands, can transduce signals from the cell interior.^23^ However, the studies show that activation of β1ARs in the Golgi apparatus from the pre-existing receptor pool is prevalent rather than by internalized receptors.^30^ Considering that many GPCR types reside in various cellular membranes^31–35^, it is likely that endomembrane signaling is a crucial component of the overall GPCR pathway, especially given that most synthetic drugs are cell permeable. Therefore, endomembrane heterotrimeric G protein signaling appears to be involved in several fatal diseases. Among these conditions, cardiovascular diseases, such as hypertrophy and heart failure are prominent.^23, 26^ Therefore, it is imperative to understand how the selective GPCR activation is regulated to develop them as therapeutic tools.

One of the major challenges in examining endomembrane signaling is the exclusive activation of endomembrane GPCRs without interferences from residual plasma membrane bound GPCR activation. Though cell-permeable ligands are available, they can activate both plasma membrane and endomembrane GPCRs. Therefore, cell impermeable antagonists are used at high concentrations to significantly inhibit the plasma membrane signaling, allowing observation of primarily endomembrane-specific signaling.^26^ However, the residual plasma membrane bound GPCR signaling can still interfere with the observations, while supersaturated levels of antagonists can also leak into the endomembranes, inhibiting the endomembrane signaling events. Therefore, achieving precise subcellular control of GPCRs to activate G proteins and control cell behavior is crucial. We have led the way by developing the first optogenetic approach for subcellular GPCR signaling control using photoreceptor opsins,^36, 37^ while achieving sub-micron subcellular resolution and sub-second ON-OFF kinetics of G protein activity in living cells.^38^ Honed by this expertise, we show herein the first-of-their-kind innovative approaches to optically activate endomembrane βARs exclusively using our photosensitive Isoproterenol-OptoIso.

## 2. RESULTS AND DISCUSSION

### 2.1 Synthesis and structure validation of protected hydroxyl Isoproterenol

**Figure 1.**
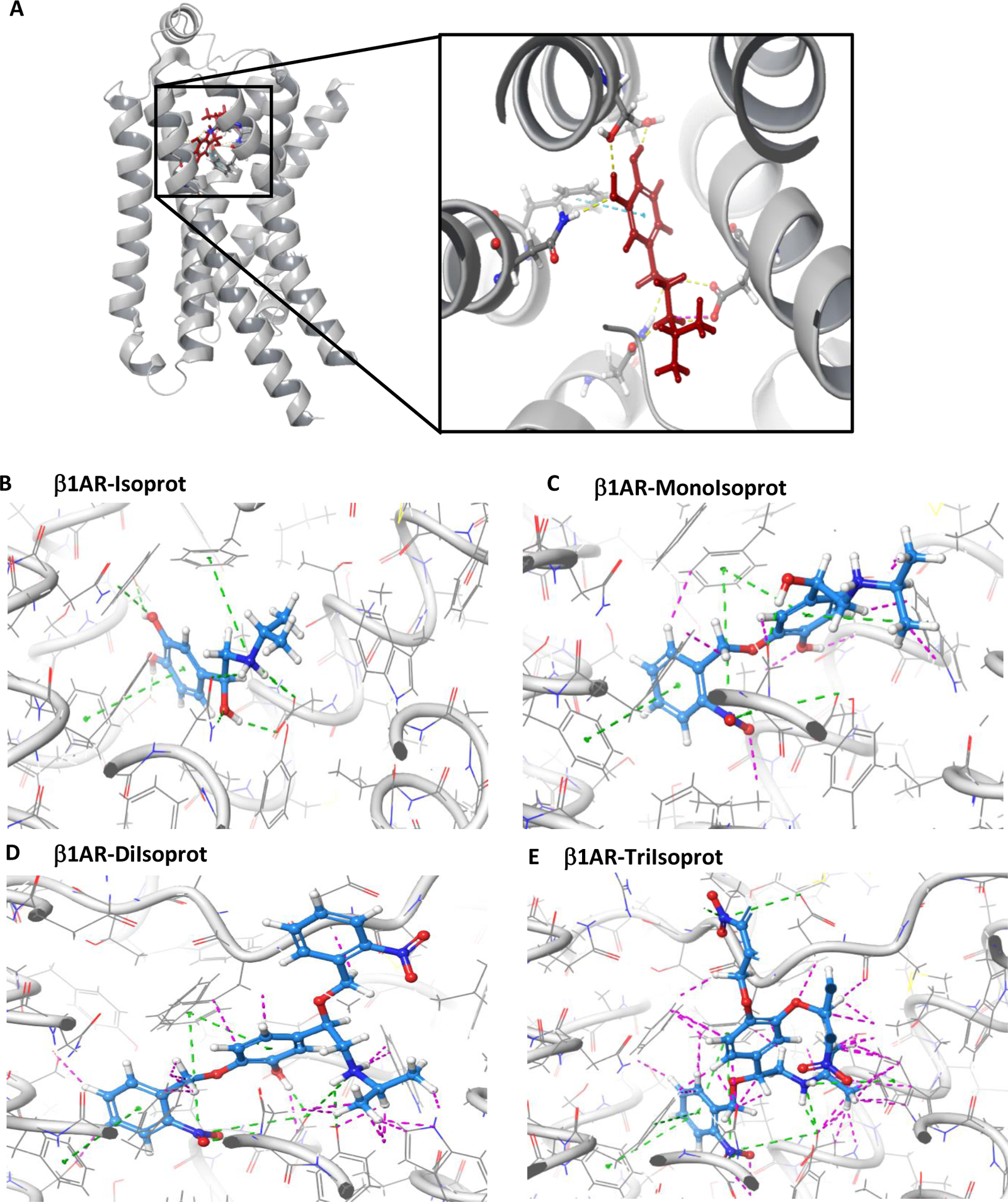
**(A)** Crystal structure of ligand-bound β1-adrenergic receptor (PDB ID: 7JJO) showing interactions with residues Ser215, Ser211, Asn310, and Asp121 in the ligand binding pocket. Yellow dotted lines: Hydrogen bonds, Blue dotted lines: π-π interactions, pink dotted lines: salt bridges. **(B)** Molecular docking of Isoproterenol to the ligand-binding pocket of the β1-adrenergic receptor (PDB ID: 7JJO) **(C)** Molecular docking of Monoprotected Isoproterenol to the ligand-binding pocket of β1-adrenergic receptor (PDB ID: 7JJO), Compared to Isoproterenol, Monoprotected Isoproterenol has more unfavorable clashes. **(D)** Molecular docking of Diprotected Isoproterenol to the ligand-binding pocket of the β1-adrenergic receptor (PDB ID: 7JJO), Compared to Isoproterenol and Monoprotected Isoproterenol, Diprotected Isoproterenol has more unfavorable clashes. **(E)** Molecular docking of Triprotected Isoproterenol to the ligand-binding pocket of the β1-adrenergic receptor (PDB ID: 7JJO), Compared to Isoproterenol, Monoprotected Isoproterenol and Diprotected Isoproterenol, Triprotected Isoproterenol has the highest unfavorable clashes. In B-E, favorable interactions are shown in green dotted lines and unfavorable clashes are shown in magenta dotted lines.

We designed several Isoproterenol (Isoprot) derivatives with the goal of discovering one with the following characteristics: (a) no activity towards β1AR before blue light exposure, (b) rapid release of the active ligand upon cell-friendly blue light exposure conditions, (c) sufficient cell permeability compared to Isoprot. Structural data of β1AR (Fig. 1A, PDB ID: 7JJ0)^39^ shows multi-point interactions between Isoprot hydroxyl and amine groups with the amino acid residues in the ligand binding cavity of the receptor.^40^ Since hydroxyl groups form key interaction with residues such as Ser215, Ser211, Asn310, and Asp121, we hypothesized that, once they are masked, the electrostatic interactions with the amine group and π-π stacking with the ligand alone would not be sufficient for ligand binding. Our goal was to create a variant of Isoprot in which the hydroxyl groups are masked using photocleavable groups.^41^ We also expected sufficient cell-permeability in the protected hydroxyl Isoprot derivatives.^42^

We began by taking Isoprot and treating it with excess 2-nitrobenzylbromide in acetonitrile in the presence of potassium carbonate as a base (Scheme 1A). Ethers of 2-nitrobenzyl are able to undergo a Norrish-type II reaction upon photolysis, releasing the free alcohols under relatively rapid irradiation conditions (Scheme 1B).^43^ The synthesis provided good chemoselectivity for OH alkylation of all three hydroxyl groups over NH alkylation. The identity of the desired product was corroborated by ^1^H and ^13^C NMR (Figure S1 A and B-before blue light). The exact mass-to-charge ratio (m/z) match for tris-*o*-nitrobenzyl triprotected Isoprot (Isoprot•NBN3-TriIsoprot) was detected by directly injecting a 1 μL of the synthesized, purified compound on the high-resolution mass spectrometry (HRMS). Verification of the structure of the TriIsoprot was done using steeped collision energy fragmentation scans (Fig. S2A and B). This experiment showed the expected fragments upon increased collision of the TriIsoprot with the nitrogen gas inside the HRMS (Fig. S2C).

With the proligand TriIsoprot in hand, we wanted to ensure that it would indeed be photolabile. The aqueous solubility of TriIsoprot was poor as judged by ^1^H NMR in D_2_O, so it was dissolved in dimethylsulfoxide-*d*_6_ (DMSO), followed by irradiation using a Kessil A160WE Tuna Blue lamp (in blue mode). During this time, the ^1^H NMR spectrum was recorded at intervals (Fig. 2). After only 15 minutes, the protected benzylic ether of TriIsoprot was observed to be completely deprotected, while a partial deprotection of the catechol ethers occurred. Meanwhile, irradiation under UV light (365 nm) gave similar results (Fig. S1A), but due to the decreased intensity (a Spectroline E-Series lamp with 4 W output was used), the deprotection took longer. Considering that the partially deprotected ligands might also give rise to a βAR partial ligand, we also prepared Isoprot•NBN_2_ (DiIsoprot) and Isoprot•NBN_1_ (MonoIsoprot) for comparison. The DiIsoprot was prepared by first synthesizing TriIsoprot, then by selectively deprotecting under blue light photolysis (the reaction was monitored by ^1^H NMR spectroscopy), followed by isolation. The MonoIsoprot was synthesized by adding one equivalent of the electrophile during synthesis, and careful isolation gave a mixture of the two mono catechol ether products.

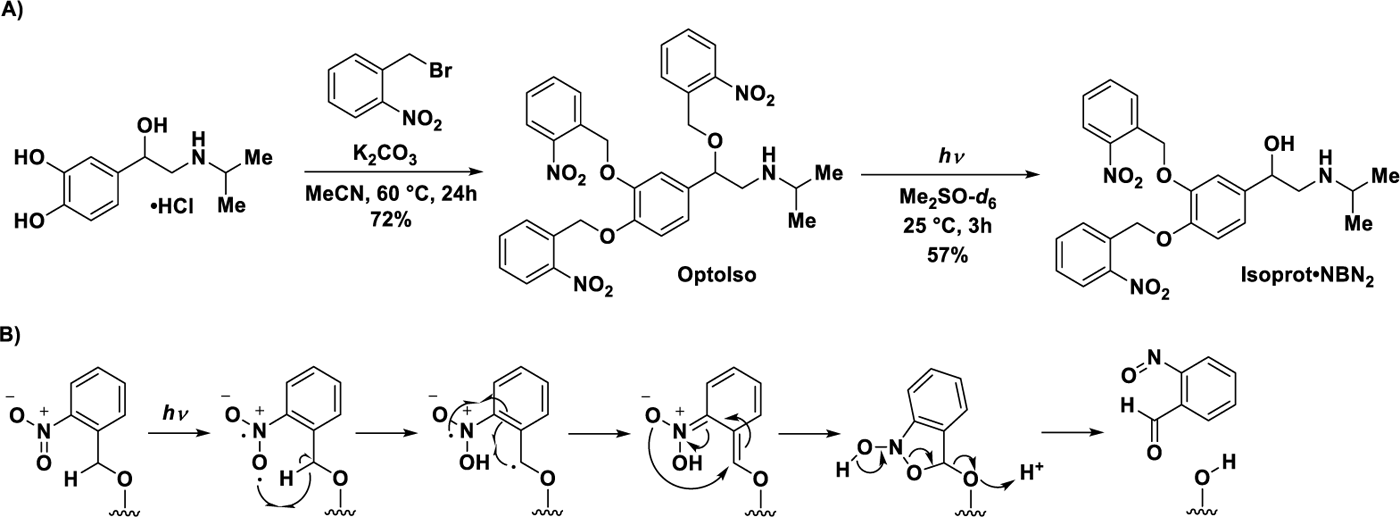

**Scheme 01**. Relevant Chemical Reactions and Mechanisms. **A.** Synthesis of Isoprot•NBN_3_ (TriIsoprot) and Isoprot•NBN_2_ (DiIsoprot) from Isoprot. **B.** Mechanism for Photolysis by a Norrish-type II Deprotection Mechanism

**Figure 2.**
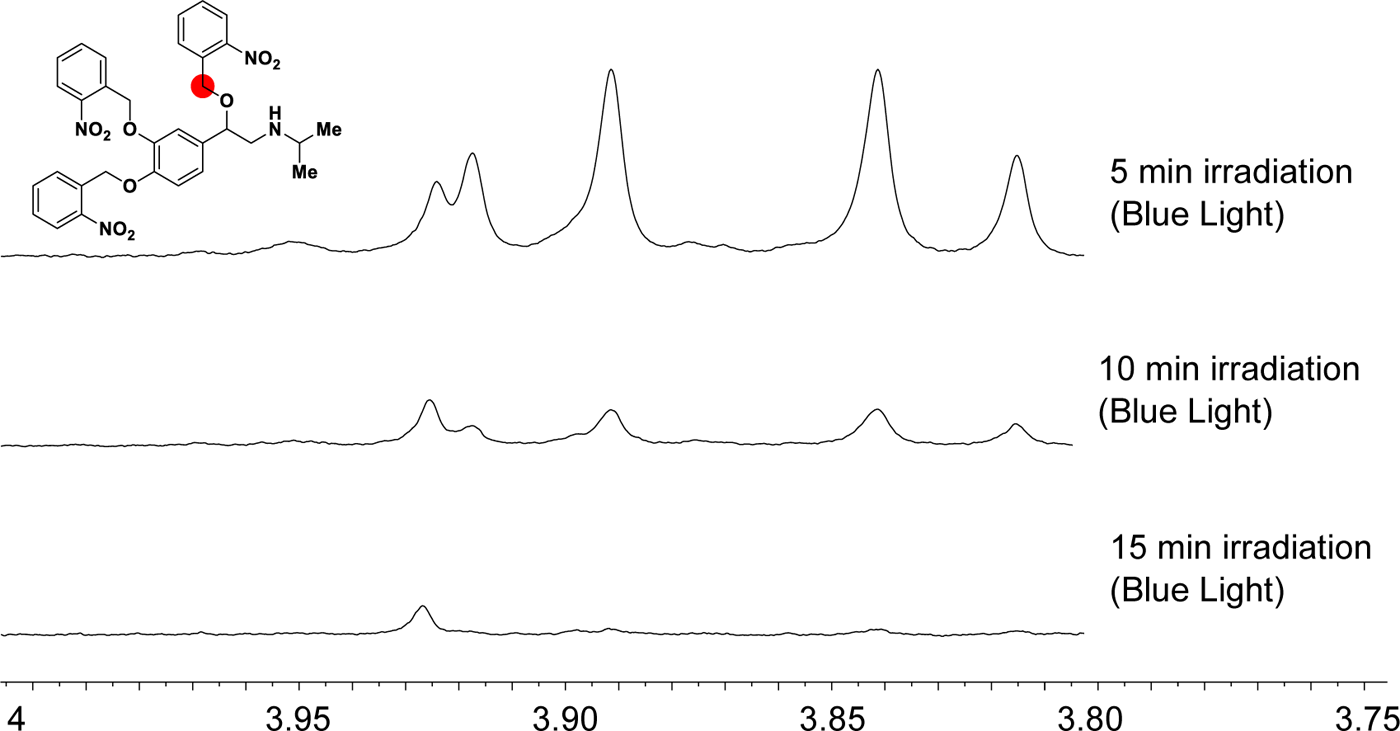
^1^H NMR Analysis of the Photodeprotection of TriIsoprot using a Tuna Blue Lamp (DMSO-*d*_6_, 600 MHz, 298 K). Spectrum has been zoomed to note the deprotection of the benzylic ether of Isoproterenol.

### 2.2 Basal and blue light-induced β1AR activation by protected Isoproterenols

We first examined the β1AR activation ability of Isoprot and its nitro benzylated variants: MonoIsoprot, DiIsoprot, and TriIsoprot in HeLa cells. Since β1AR is a Gs-coupled GPCR, we used Venus-miniGs recruitment to the plasma membrane to examine receptor activation.^44^ In HeLa cells expressing β1AR-CFP and Venus-miniGs, we examined Venus-miniGs dynamics upon the addition of pre-blue light exposed Isoprot or its analogs to measure their activities using time-lapse imaging. For imaging, cells with moderate fluorescence expression with pixel intensities ranging from 200,000 to 400,000 at ∼6 µW, and 40 ms exposure were selected for all the experiments. We used the reduction of cytosolic Venus fluorescence due to miniGs plasma membrane recruitment as the indicator of β1AR activation. When the fluorescence intensity differences before the ligand addition and at the equilibrium were considered, the highest Venus-miniGs recruitment was observed in cells exposed to Isoprot, while partial responses were observed in cells exposed to MonoIsoprot and DiIsoprot (Fig. 3A-cell images and the plot). TriIsoprot did not show any probe recruitment (Fig. 3A-bottom images, plot-blue curve). The data indicated that, although MonoIsoprot and DiIsoprot induced significant basal activity, TriIsoprot did not induce a measurable activity. Next, we examined whether these compounds could activate β1AR upon photo-deprotection using blue light. We exposed 100 μM Isoprot, or its protected analogs, to blue light (405 nm, 3.28 mW/mm^2^, 1 cm away) for 15 minutes in DMSO and diluted to have 500 µM solutions in cell culture media. We then added these solutions to HeLa cells expressing the same constructs to have a final ligand concentration of 100 μM. All compounds, including TriIsoprot showed Venus-miniGs recruitment (Fig. 3B-cell images). We measured the relative light sensitivity of each compound by measuring the baseline normalized extents of Venus-miniGs loss in the cytosol due to plasma membrane recruitment and compared the values without and with blue light exposure conditions. Since Isoprot has no photolabile groups, it showed similar responses upon the addition of compounds with and without light exposure. The analysis captured this by showing near-zero relative light sensitivity (Fig. 3C). TriIsoprot showed the highest light sensitivity in this assay.

To confirm the exclusive β1AR activation ability of the above four compounds, we also examined nanobody-80 (Nb80) recruitment from the cytosol to the plasma membrane in HeLa cells expressing Nb80-mCherry and β1AR-CFP following a procedure similar to the above miniGs experiment. Nb80 exclusively binds active state β1ARs.^26^ The data show that Nb80 responses also followed an analogous behavior to miniGs, in which TriIsoprot did not show a detectable basal activity before and exhibited a robust Nb80 recruitment after 15 min blue light exposure of the compound (Fig. S3).

**Figure 3.**
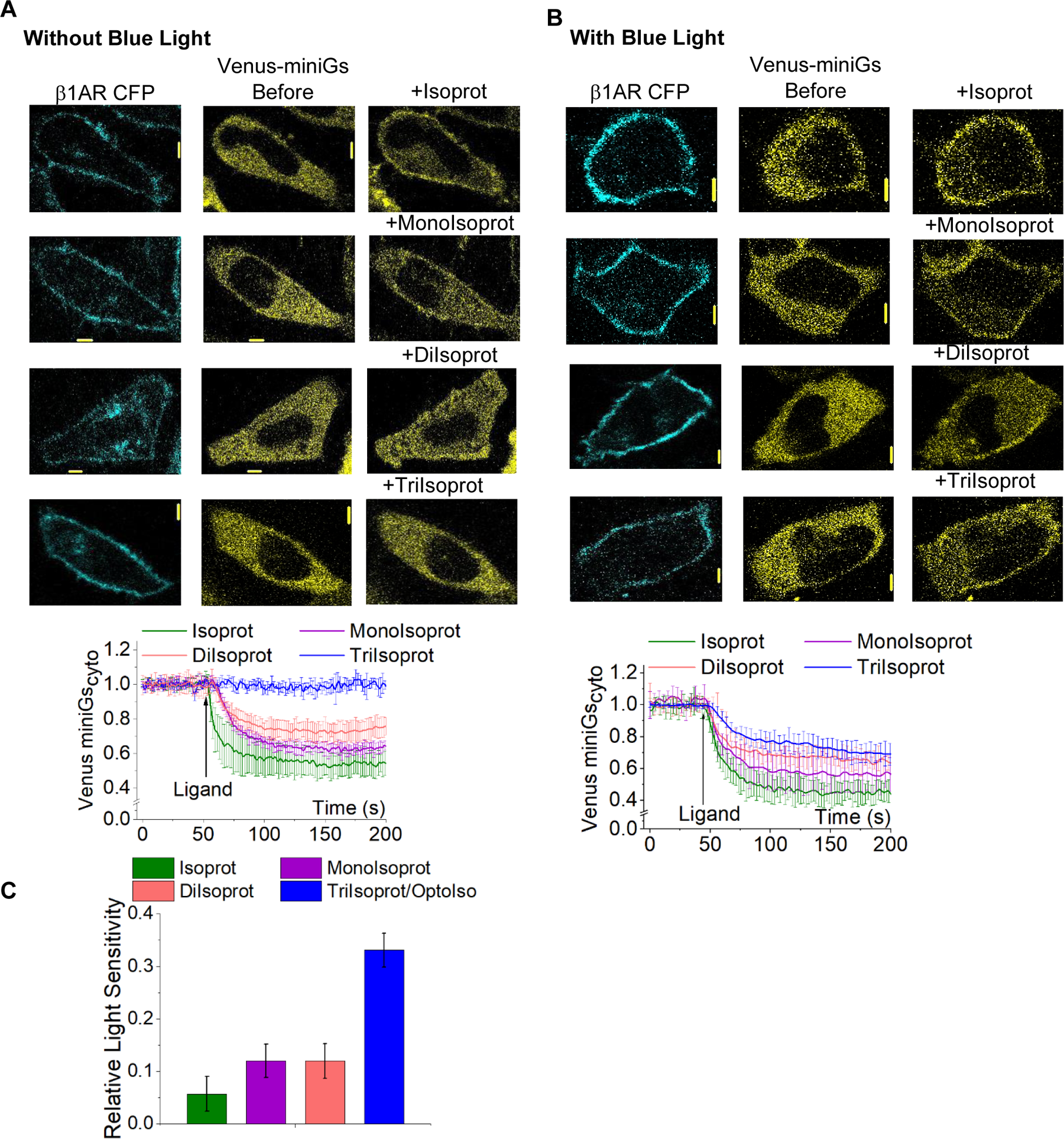
**(A)** HeLa cells expressing β1AR-CFP and Venus-miniGs were treated either with 100 μM Isoproterenol (Isoprot) or three protected isoproterenol variants; Mono protected (MonoIsoprot), diprotected (DiIsoprot) and tri-protected (TriIsoprot/OptoIso) at 50 seconds (all the compounds are not treated with blue light). Isoprot-treated cells showed robust Venus-miniGs recruitment to the plasma membrane and MonoIsoprot and DiIsoprot-treated cells showed a detectable miniGs translocation to the plasma membrane. TriIsoprot-treated cells didn’t show a detectable Venus-miniGs translocation to the plasma membrane of the cells. The line plot shows the cytosolic Venus-miniGs dynamics normalized to the basal level fluorescence. **(B)** HeLa cells expressing β1AR-CFP and Venus-miniGs were treated either with 100 μM of Isoprot or three protected isoprot variants; MonoIsoprot, DiIsoprot and OptoIso at 50 seconds, after exposing the compounds to blue light for 15 minutes. Isoprot-treated cells showed a robust Venus-miniGs recruitment to the plasma membrane and MonoIsoprot and DiIsoprot-treated cells showed a detectable MiniGs translocation to the plasma membrane. OptoIso-treated cells also showed significant Venus-miniGs translocation to the plasma membranes of the cells. The line plot shows the cytosolic Venus-miniGs dynamics normalized to the basal level fluorescence. **(C)** The bar chart represents the light sensitivity of the compounds. Here, the difference in average sensor recruitment extents before and after blue light exposure was plotted as a measurement of light sensitivity. The OptoIso showed the highest light sensitivity and Isoprot showed no light sensitivity. Average curves were plotted using cells from 4 independent experiments. The error bars represent SD (standard deviation of mean). The scale bar = 5 µm. CFP: Cyan Fluorescent Protein; MonoIsoprot: Monoprotected Isoproterenol; DiIsoprot: Di-protected Isoproterenol; OptoIso: Tri-protected Isoproterenol; Cyto: cytosolic.

Since TriIsoprot does not show any activity before blue light exposure and brings β1AR to the active conformation upon blue light exposure, we named it “OptoIso” and henceforth used to optically activate β1AR in living cells.

To assess the β1AR activation efficacies of the four compounds, we expressed β1AR-CFP and Venus-miniGs in HeLa cells and treated the cells with different concentrations (10 nM to 1 mM) of each compound pre-exposed to blue light for 15 minutes as described before. We added 10 nM of blue light exposed OptoIso at t=30 seconds after the onset of YFP imaging and continued until 180 seconds, at which the cytosolic fluorescence reached equilibrium. At this point, we added the next higher concentration (100 nM) of blue light exposed OptoIso. We repeated this procedure for the same cell culture dish for 1 μM, 10 μM, 100 μM, 1 mM, and 10 mM concentrations (Fig. 4A - cell images). Using the equilibrium responses of Venus-miniGs after each addition, we generated dose-response curves for OptoIso and blue-light-exposed Isoprot, MonoIsoprot, and DiIsoprot (Fig. 4A - plot). Using these dose-response curves, we calculated the EC50 values.

Compared to the ∼50 nM EC50 of Isoprot previously reported by examining the inhibition of contractions in isolated, field stimulated rat vas deferens^45^, the EC50 of Isoprot we observed (250 nM) is 5-fold higher. Nevertheless, given that we calculated EC50 value using single-cell data based on receptor conformational change observed in live cells, we believe our analysis is an accurate measure of β1AR activation efficacy. The same assay for 15 min-blue light exposed OptoIso showed 81 μM EC50. MonoIsoprot and DiIsoprot showed 0.63 μM and 20 µM EC50, respectively. These data indicated that upon blue light exposure, OptoIso is likely to undergo partial deprotection. This observation is validated by our ESI-MS data showing m/z= 617 for OptoIso before blue light exposure and generation of m/z=482 (DiIsoprot) and 346 (MonoIsoprot), in addition to the remaining OptoIso+H^+^ indicated by m/z= 617 signal (Fig. S1B). Also, based on the blue light deprotection of MonoIsoprot and DiIsoprot, it is likely that, upon blue light exposure, the majority of OptoIso becomes mono-deprotected, generating DiIsoprot, while only a fraction is converted to MonoIsoprot. This should not be misconstrued as a result of inefficient blue light deprotection. When cells are in culture media containing 100 µM OptoIso, the efficient miniGs recruitment with 200 second t_1/2_ upon blue light exposure shows that the underlying photo-deprotection is an efficient process (Fig. S3C). We also found that continuous blue light exposure of OptoIso in DMSO beyond 15 minutes resulted in less efficient β1AR activation (Fig. S3D).

**Figure 4.**
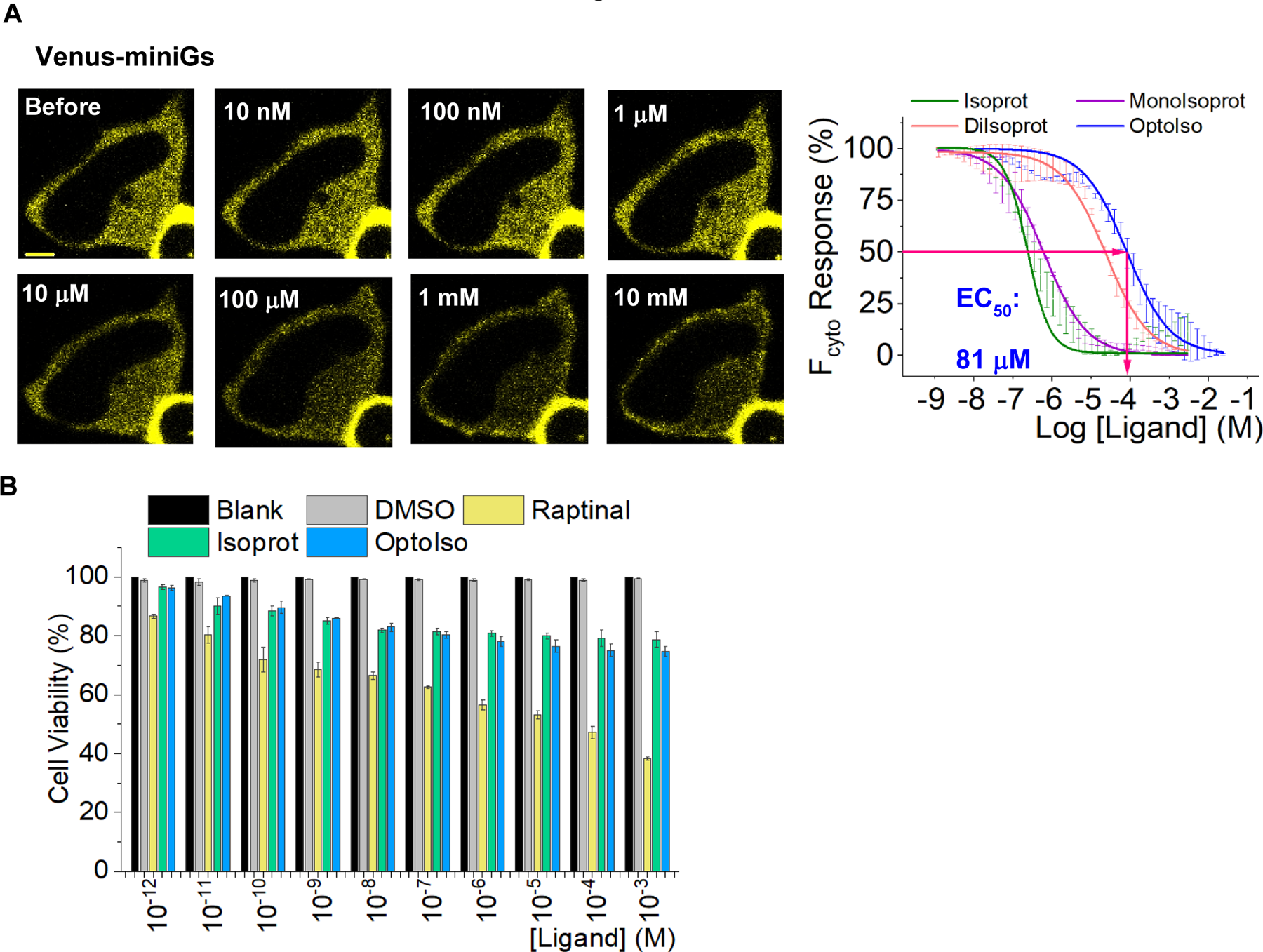
**(A)** Cell images and EC_50_ curves generated with Venus-miniGs response for different ligand concentrations. Cell images show a cumulative effect from the addition of the different concentrations of the ligands. When the concentration of the ligands was increased the sensor recruitment to the activated receptor increased subsequently. The plot was generated with the sensor recruitment extent in different concentrations normalized from 0 to 100. Then the data was fitted to the pharmacology category dose response algorithm under nonlinear curve fitting in Origin Pro software. The scale bar = 5 µm. **(B)** The bar chart represents the cell viability (%) determined by MTT assay after 1 hour incubation followed by drug treatment. The data shows no significant cytotoxicity from the modified ligands compared to the controls. Average cell viability (%) was plotted using cells from 3 independent experiments. The error bars represent SD (standard deviation of mean). MonoIsoprot: Monoprotected Isoproterenol; DiIsoprot: Diprotected Isoproterenol; OptoIso: Triprotected Isoproterenol

We further examined whether the OptoIso concentrations we used induced significant cytotoxicity by measuring the cell viability using the MTT assay.^46^ First, we first plated HeLa cells on a 96-well cell culture plate, and the next day, we incubated them with different concentrations (1 pM to 1 mM) of OptoIso for 1 hour. Next, we washed the cells and incubated them with a 5% MTT solution for 1 hour, followed by washing with PBS and then incubating cells with 1 µL DMSO for 1 hour before measuring the absorbance at 540 nm. As a positive control, we exposed cells to raptinal, a compound known to induce apoptosis,^47^ for 1 hour. Interestingly, untreated and DMSO-treated controls did not show significant cytotoxicity (Fig. 4B-black and gray bars in the plot), while Isoprot and OptoIso-treated cells showed similar cytotoxicity (Fig. 4B-green and blue bars in the plot). We found that the cell viability of raptinal-treated cells was lower than that of Isoprot and OptoIso-treated cells, indicating significant cytotoxicity as expected (Fig. 4B -yellow bars in the plot).

### 2.3 OptoIso allows for reversible optical control of β1AR

We and others have used photopigment GPCRs such as rhodopsin and color opsins to activate G protein signaling upon reversible light stimulations.^48, 49^ Opsin-mediated signaling is fast and can be precisely controlled in diffraction-limited sub-micrometer regions of interest in single cells using confined light stimuli.^36, 50, 51^ The resultant activity can be reversed immediately upon the termination of light stimulation.^48–51^ Although photocleavable small molecules has been used to activate cell surface GPCRs optically,^52, 53^ to our knowledge, such methods did not allow reversible control. As predicted by the Stokes–Einstein equation, studies show that small molecules, including salbutamol (a βAR ligand), diffuse significantly faster (∼6.6×10^6^ cm^2^/s) in aqueous media.^54^ If we momentarily photoconvert OptoIso to generate ∼1 µM βAR-activating form in a limited media volume of ∼1.5 µL next to a cell in a dish with 1 mL culture media, calculations show that the ligand concertation in the bulk media to be ∼15 nM seconds after blue light termination. This indicated the possibility of using OptoIso to reversibly switch on and off βARs on blue light command.

To examine OptoIso-β1AR reversible activation in living cells, we expressed β1AR-CFP, Venus-miniGs, and Nb80-mCherry in HeLa cells and added 100 μM OptoIso to the cell culture media. We then exposed selected cells to blue light (405 nm, 3.28 mW/mm^2^), starting from t=60 to 480 seconds (Fig. 5A-1^st^ activation-blue light, plot B and C first blue box). During this process, we collected time-lapse images of cells for Venus (515 nm ex/ 540 nm em, 12.3 µW/ mm^2^) and mCherry (594 nm ex/ 620 nm em, 9.7 µW/ mm^2^) at 2-second intervals. Blue light exposure stimulated a gradual yet robust Venus-miniGs and Nb80-mCherry recruitment to the plasma membrane, indicating the β1AR activation. Then, we terminated the blue light and continued imaging the sensors to allow for diffusion of the photoproducts into the bulk media. Over time, both of the sensors translocated back to the cytosol, indicating the recovery of the β1AR activity with recovery t_1/2_ of ∼14 minutes for miniGs and ∼17 minutes for Nb80 (Fig. 5A-Recovery-No Blue light, plot B and C no Blue light region). As a control, we then re-initiated the blue light to make sure that the observed sensor reversal was not due to β1AR desensitization. Indicating that, indeed, the recovery was due to ligand diffusion and β1AR deactivation, blue light induced the re-recruitment of the sensors to the plasma membrane (Fig. 5A-2^nd^ activation-blue light, plot B and C second blue box). Considering the likely fast diffusion of the βAR ligand generated through photo uncaging, the somewhat slow recovery rates of the sensors indicate that the recovery process is limited by the ligand dissociation from the β1AR. Collectively, these data clearly show that OptoIso allows for reversible optical control of βARs (Movie S1 and S2).

**Figure 5.**
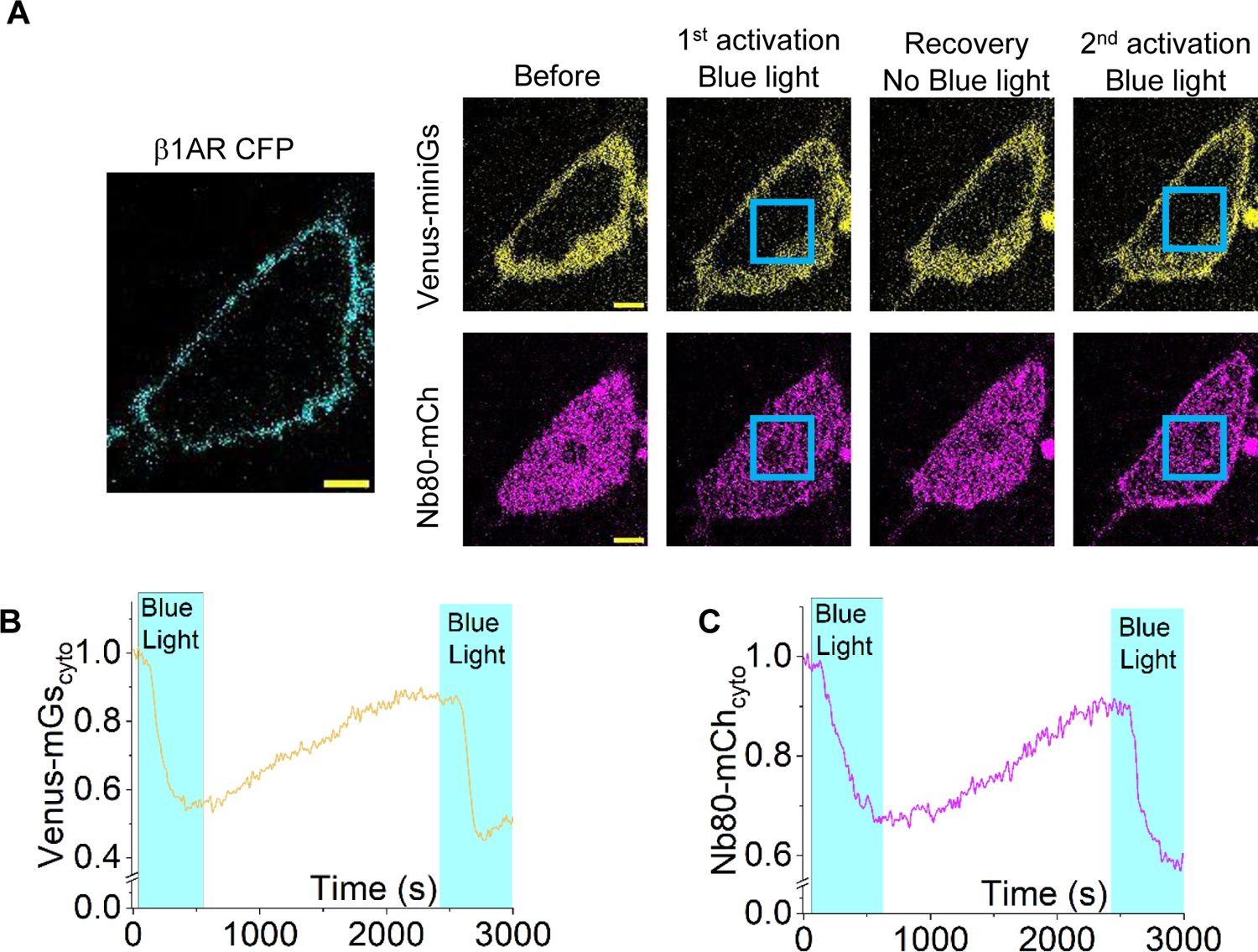
**(A)** HeLa cells expressing β1AR-CFP, Venus-miniGs, and Nanobody80-mCherry were treated with 100 mM OptoIso. Blue light exposure was started at 1 minute and the sensors moved to the plasma membrane indicating the receptor activation. Then the blue light was terminated allowing the sensors to recover. After that cells were exposed to blue light again and the sensor recruitment to the plasma membrane indicated a second activation of the receptors. **(B)** The line plot shows the cytosolic Venus-miniGs dynamics normalized to the basal level fluorescence. **(C)** The line plot shows the cytosolic Nanobody80-mCherry dynamics normalized to the basal level fluorescence. Both line plots show the activation upon blue light exposure, reversibility with the termination of the blue light exposure, and a subsequent activation with a second blue light exposure. Single cell data were plotted after repeating for 3 independent experiments.

### 2.4 OptoIso allows for optical activation of β1AR downstream signaling

Upon GPCR activation, Gα exchanges the bound GDP to GTP, dissociating the heterotrimer into Gα_GTP_ and Gβγ.^55^ Following the dissociation, Gγ members reversibly translocate from the plasma membrane to the endomembranes as Gβγ, and the process is Gγ-type dependent.^56^

**Figure 6.**
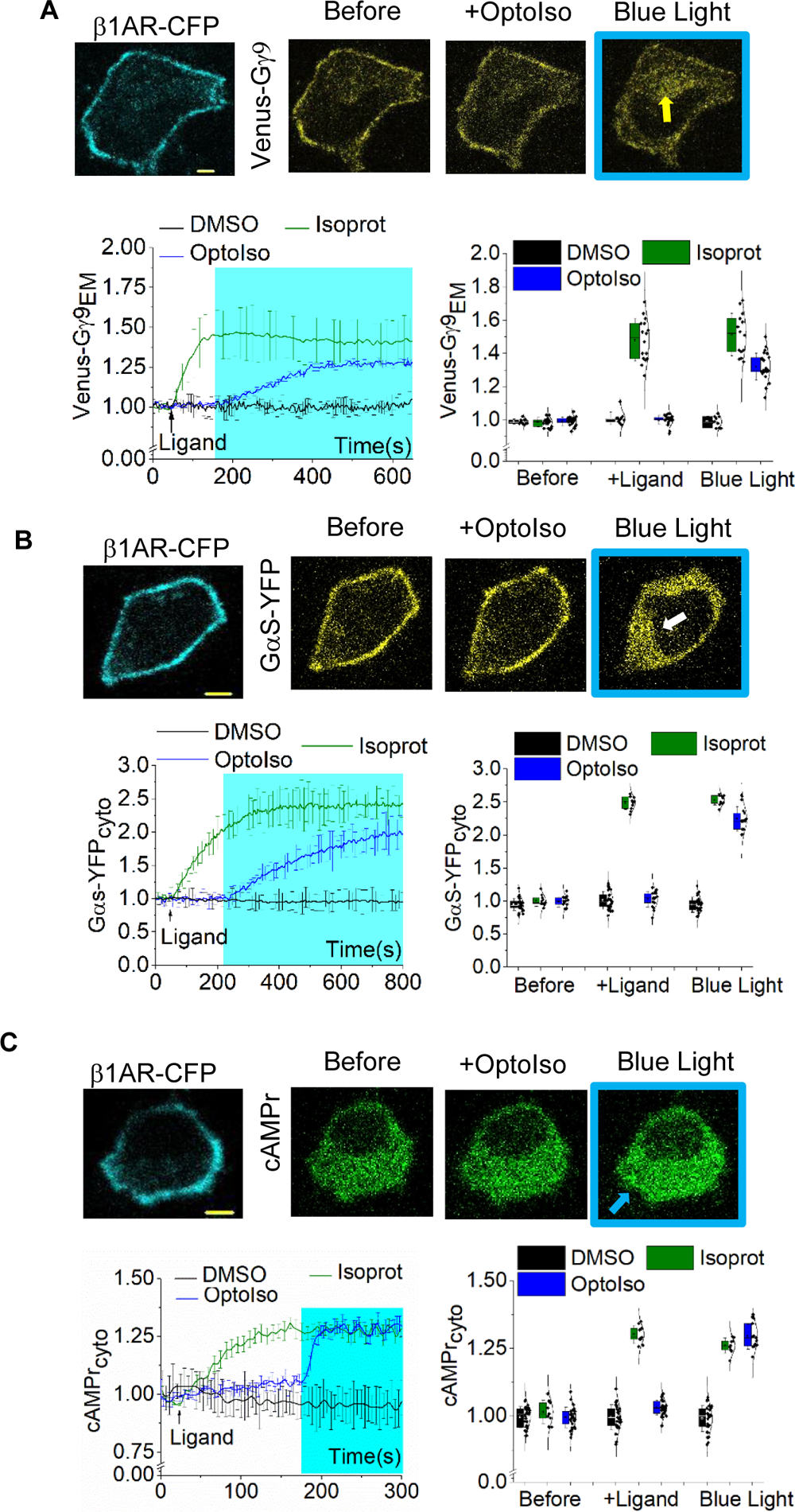
(A) HeLa cells expressing β1AR-CFP and Venus-Gγ9 was treated with 100 μM OptoIso at 50 seconds and blue light was given at 3 minutes. The cells showed significant Gβγ translocation upon exposure to blue light. The line plot and the whisker box plot show the endomembrane Venus-Gγ9 dynamics normalized to the basal level fluorescence under different conditions. The yellow arrow indicates the endomembrane accumulation of the fluorescence protein. **(B)** HeLa cells expressing β1AR-CFP and Gαs-YFP were treated with 100 μM OptoIso at 50 seconds and blue light was given at 3 minutes. The cells showed robust Gαs translocation upon exposure to blue light. The line plot and the whisker box plot show the cytosolic Gαs-YFP dynamics normalized to the basal level fluorescence under different conditions. The white-colored arrow indicates the cytosolic fluorescence increase. **(C)** HeLa cells expressing β1AR-CFP and cAMPr (cAMP sensor) were treated with 100 μM OptoIso at 50 seconds and blue light was given at 3 minutes. The cells showed robust cAMPr fluorescence increase upon exposure to blue light. The blue-colored arrow indicates the whole cell fluorescence increase. The line plot and the whisker box plot show the cytosolic cAMPr dynamics normalized to the basal level fluorescence under different conditions. Average curves in each experiment were plotted using cells from 4 independent experiments. The error bars represent SD (standard deviation of mean). The scale bar = 5 µm. CFP: Cyan Fluorescent Protein; Isoprot: isoproterenol; OptoIso: Tri protected Isoproterenol; Cyto: cytosolic; EM: endomembrane; DMSO: Dimethyl Sulfoxide. The blue box indicates the blue light exposure.

Using Gγ9, the Gγ subunit from the cone photoreceptor cells that provided the fastest translocation ability to Gβγ, we have measured heterotrimer activation dynamics in single cells and sub plasma membrane regions.^38^ To examine OptoIso-activated β1AR signaling in living cells, we exposed HeLa cells expressing Venus-Gγ9 and β1AR-CFP to 100 μM OptoIso at t=50 seconds (after the onset of time-lapse imaging) and to blue light (405 nm, 3.28 mW/mm^2^) from t=180 to 600 seconds (Fig. 6A). During this process, we collected time-lapse images of cells for Venus (515 nm ex/ 540 nm em, 12.3 µW/ mm^2^) at 1 second intervals. Blue light exposure stimulated a gradual yet robust Venus-Gγ9 translocation (Fig. 6A, yellow arrow). In a similar experiment, when we exposed cells to 1μL of DMSO, regardless of the blue light exposure, the cells did not show a detectable Gβγ translocation (Fig. S4A-top raw). We also performed a positive control experiment using 100 μM of Isoprot, which induced a robust Gγ9 translocation immediately upon addition, independent of blue light exposure (Fig. S4A-bottom raw).

It has been shown that upon β1AR stimulation, Gαs translocates from the plasma membrane to the cytosol, suggesting that the process occurs through lipid rafts.^57^ To examine Gαs heterotrimer activation by blue light-exposed OptoIso, we imaged Gαs-YFP (a circularly permuted Gαs)^58^ dynamics in HeLa cells expressing β1AR-CFP (Fig. 6B). We performed time-lapse imaging of cells for YFP (1Hz) and added OptoIso (100 μM) at t=50 seconds. From t=180 to 800 seconds, cells were exposed to blue light while capturing YFP time-lapse images. Although OptoIso addition did not induce a detectable change in cytosolic YFP fluorescence, blue light exposure induced a gradual and robust increase due to Gαs-YFP translocation (Fig. 6B-white arrow). As a negative control, we performed a similar experiment using the vehicle solvent (DMSO) and the cells did not show Gαs -YFP translocation (Fig. S4B-top). As a positive control, addition of Isoprot (100 μM) at t=50 seconds induced a fast, blue light independent Gαs-YFP translocation (Fig. S4B-bottom).

We next examined the β1AR activation-induced adenylyl cyclase stimulation and cAMP generation in HeLa cells expressing β1AR-CFP and the cAMP sensor, cAMPr.^59, 60,61^ Similar to the Gαs experiment above, we added 100 μM of OptoIso at t=50 seconds, and exposed cells to blue light from t=180 to 350 seconds. Cells exhibited an increase in cAMPr fluorescence only upon blue light exposure (Fig. 6C-blue arrow). The cells treated with Isoprot showed instantaneous fluorescence increase upon ligand addition (positive control-S4C-bottom). The cAMPr fluorescence in cells treated with DMSO (negative control) remained unchanged regardless of DMSO addition or blue light exposure (Fig.S4C-top). These data collectively indicated that OptoIso can be used to optically control β1AR signaling.

### 2.5 OptoIso induced endomembrane exclusive β1AR signaling

To examine the endomembrane exclusive β1AR signaling activated by cell-penetrated and blue light-activated OptoIso in HeLa cells, we first incubated cells expressing β1AR-CFP, Venus-miniGs, and Nb80-mCherry with 100 μM of OptoIso for 30 minutes. We next washed the cells with Hank’s Balanced Salt Solution (HBSS) 5 times to remove excess extracellular OptoIso before starting imaging in cell culture media. Upon exposing cells to blue light starting at t=50 seconds, we observed a robust Venus-miniGs and Nb80-mCherry recruitment to the endomembrane regions with prominent β1AR-CFP expression (Fig. 7A, yellow arrows). To validate that the observed endomembrane miniGs and Nb80 recruitments are due to β1AR activation, we next exposed cells to the cell-permeable β1AR antagonist, 10 μM of metoprolol (t=720 seconds). A fast dissociation of the probes with t_1/2_ = 68 seconds for Nb80-mCherry and t_1/2_ = 46 seconds for Venus-miniGs was observed, indicating that the OptoIso-blue light activated endomembrane β1ARs. Additionally, to ensure that observed probe recruitments are not due to an artifact because of blue light exposure, we performed an analogous experiment using DMSO in place of OptoIso. As expected, blue light exposure induced neither miniGs nor nanobody80 recruitment to the endomembranes (Fig. S5A).

The traditional methods of detecting endomembrane β1AR employ subsequent exposure of cells to a β1AR antagonist with less cell permeability, followed by the addition of a cell-permeable ligand. We performed this experiment by exposing cells to 100 μM sotalol (the cell impermeable antagonist) and, next to 10 μM, dobutamine (cell-permeable β1AR agonist) (Fig. S5B).^26^ Though cells showed primarily an endomembrane probe recruitment (Fig. S5B-white arrow), since the receptor activity is governed by the relative affinities of the antagonist and the agonist, it is difficult to ensure that there is no residual signaling from the cell surface β1AR.

We next examined the ability of spatially constrained β1AR activation only to user-defined sub-endomembrane regions of selected cells in a population of cells. We prepared HeLa cells similar to the above OptoIso-blue light experiment. Upon irradiation of blue light near an endomembrane region of cell1 starting from 30 seconds, we observed Venus-miniGs and Nb80-mCherry recruitment exclusively to the adjacent sub-endomembrane region (Fig. 7B). Starting at 120 seconds, a similar area in the Cell2 was exposed to blue light and a similar Venus-miniGs and Nb80-mCherry recruitment to the adjacent sub-endomembrane was observed (Fig. 7B). These data show the feasibility of using OptoIso to trigger βAR signaling in user-defined subcellular regions in single cells using blue light while probing signaling using other wavelengths (Movie S3 and S4).

**Figure 7.**
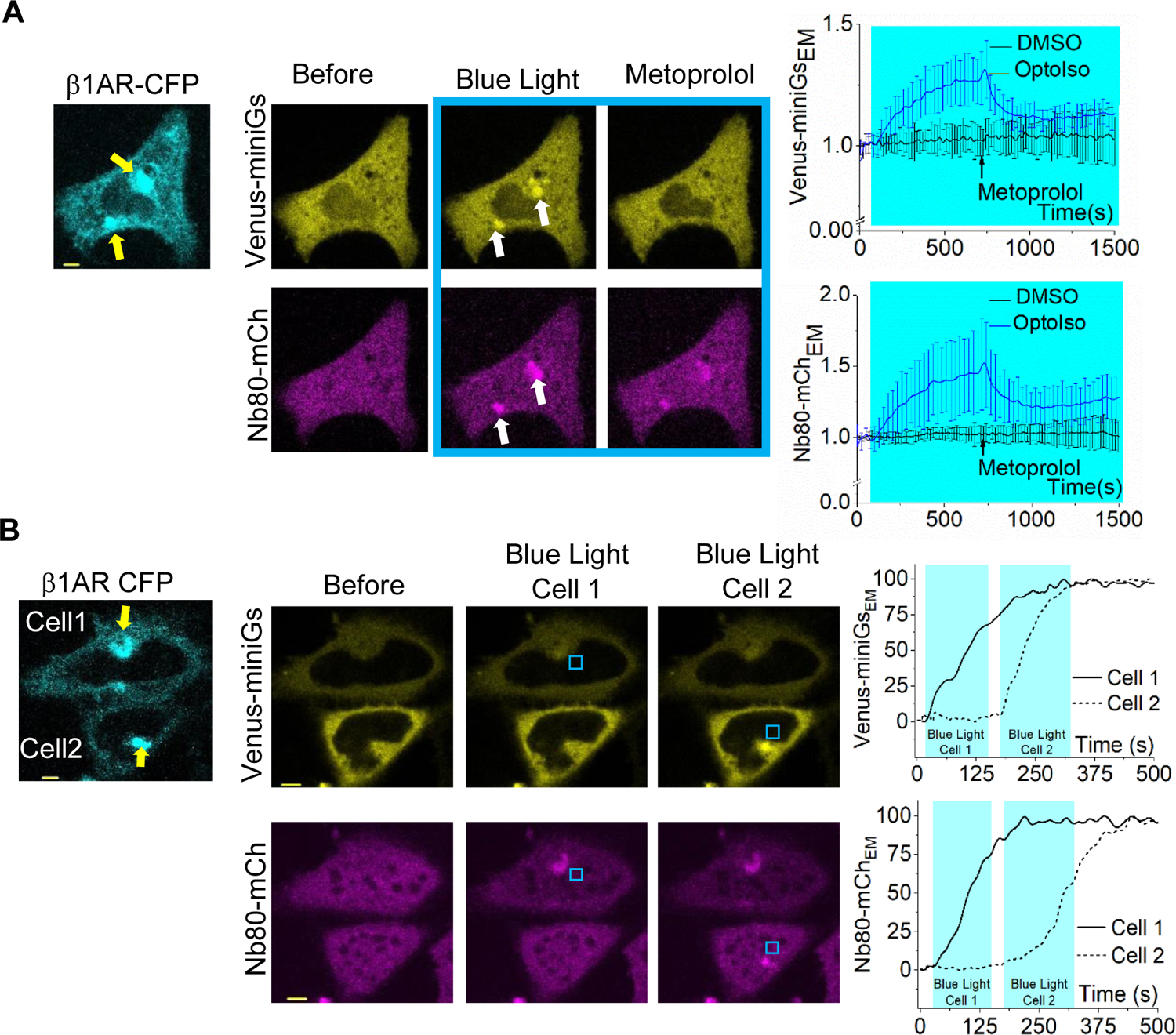
**(A)** HeLa cells expressing β1AR-CFP, Venus-miniGs and Nanobody80-mCherry were treated with 100 μM OptoIso and incubated for 30 minutes. Then the cell culture dish was washed with HBSS 5 times, and the cells were exposed to blue light at 50 seconds while imaging. A robust Venus-miniGs and Nanobody80-mCherry recruitment were observed exclusively in the endomembranes inside the cells. The disappearance of these sensors from the endomembranes was observed at 12 minutes, upon the addition of 10 μM Metoprolol. The line plots show the Venus-MiniGs and Nanobody80-mCherry dynamics at endomembranes. **(C)** HeLa cells expressing β1AR-CFP, Venus-miniGs, and Nanobody80-mCherry were treated with 100 μM OptoIso and incubated for 30 minutes. Then the cell culture dish was washed with HBSS 5 times. The blue light was exposed near the endomembrane receptors only in one cell. Venus-miniGs and Nanobody80-mCherry were recruited to the endomembranes only in that cell. Then the adjacent cell endomembranes were optically activated after 2 minutes and both sensors were recruited to the endomembranes of the respective cell. The line plots show the Venus-miniGs and Nanobody80-mCherry dynamics at endomembranes. The yellow arrows indicate the endomembrane bound receptor expression and the white arrows indicate the sensors recruitment to the endomembranes. Average curves in each experiment were plotted using cells from 3 independent experiments. The error bars represent SD (standard deviation of mean). The scale bar = 5 µm. CFP: Cyan Fluorescent Protein; OptoIso: Tri protected Isoproterenol; EM: endomembrane; DMSO: Dimethyl Sulfoxide. The blue box indicates the blue light exposure.

Although the receptor conformational changes and downstream signaling of activated β1ARs at endomembranes are investigated,^26, 27^ to our knowledge, direct evidence for G protein heterotrimer activation at endomembranes is lacking. Here, we developed an assay based on the mobility differences of Gβγ in the heterotrimer (Gαβγ), and the free Gβγ liberated upon GPCR activation. G proteins heterotrimers contain 2-3 lipid modifications: myristoyl or palmitoyl, or both on the N-terminus of Gα and prenyl modification on the C-terminus of Gγ.^62^ Therefore, mobility of heterotrimers through the cytosol is not favored, and the lateral diffusion through the membranes is a relatively slow process. On the contrary, free Gβγ shuttles efficiently through the cytosol in a Gγ subtype dependent manner, in which Gγ9 provided the Gβγ the fastest shuttling rate.^63^ To validate the Gβγ mobility assay, we examined pre and post cell surface β1AR activation induced mobility of Venus-Gγ9 by measuring Fluorescence Recovery After Half-cell Photobleaching (FRAP-hc) in HeLa cells expressing β1AR-CFP, Venus-Gγ9 and Nb80-mCherry. Briefly, using a 515 nm laser-assigned FRAP-PA unit, we exclusively photobleached half-cell Venus fluorescence in HeLa cells expressing Venus-Gγ9 and β1AR-CFP, before and after β1AR stimulation with 100 µM Isoprot (Fig. S5C). Data show that, compared to 150 seconds of Gγ9 mobility half time (t_1/2_) in cells before β1AR activation, cells exhibited 55 seconds t_1/2_ in cells with activated β1AR (Fig. S5C-whisker box plot). One-way ANOVA showed that the mobility t_1/2_ of Gγ9 before the β1AR activation is significantly higher than that of the after activation (One-way ANOVA: *F_1,38_* = 67.9416, *p* = <0.0001, Table S1A and B). This is an unprecedented 3-fold increase due to the Gβγ liberation from the heterotrimer.

**Figure 8.**
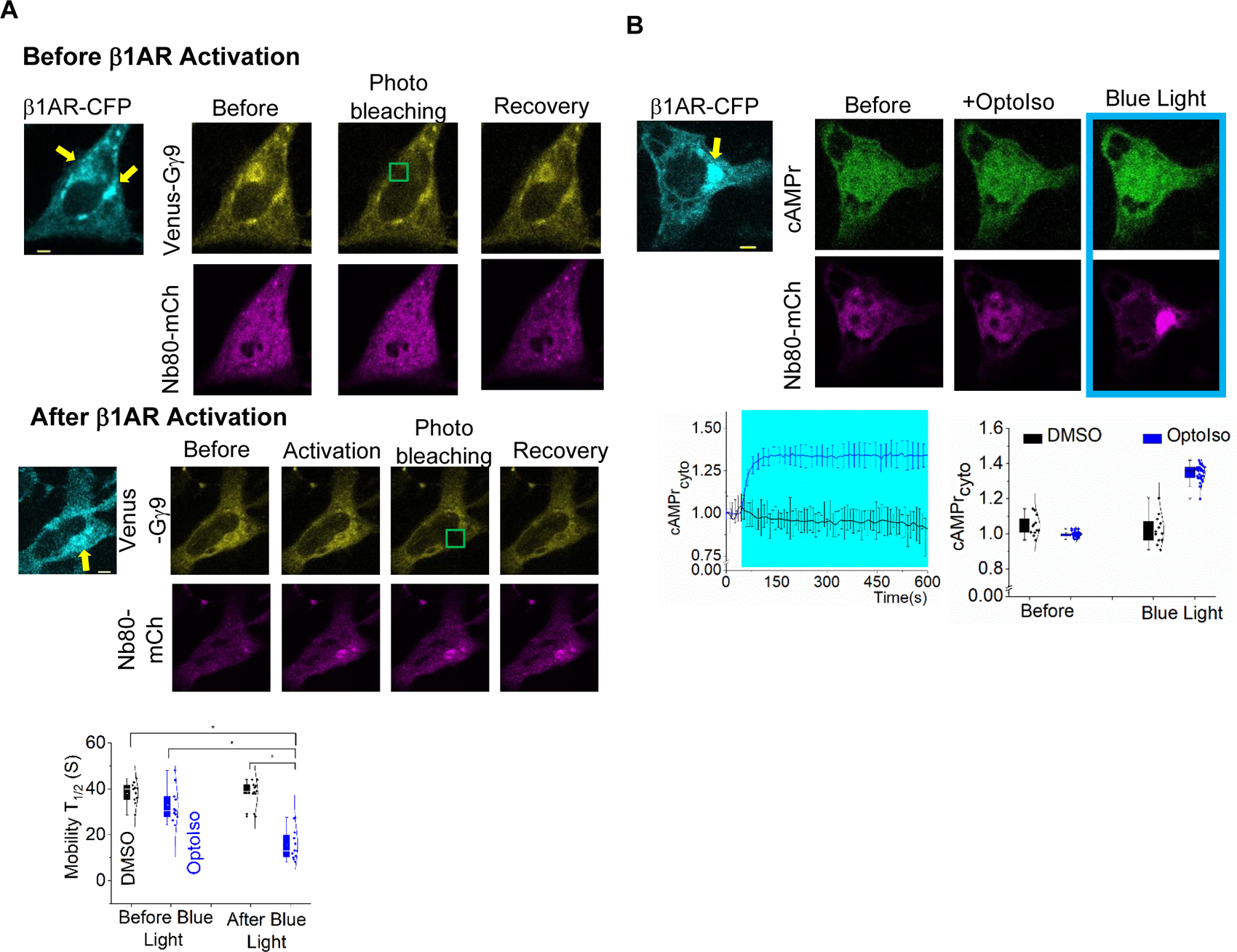
**(A)** HeLa cells expressing β1AR-CFP, Venus-Gγ9, and Nanobody80-mCherry were treated with 100 μM OptoIso and incubated for 30 minutes. Then the cell culture dish was washed with HBSS 5 times. Venus-Gγ9 at the endomembranes were photobleached and the fluorescence recovery after photobleaching was examined before the receptor activation. Nb80-mCherry was cytosolic since the receptors were not activated. Then the cells were exposed to blue light until Nb80-mCherry recruitment was observed at endomembranes showing the activation of the receptors. After that Venus-Gγ9 at the endomembranes were photobleached and the fluorescence recovery after photobleaching was examined. The whisker box plots show mobility half-time (t_1/2_) of Venus-Gγ9 before and after the blue light exposure in cells treated with DMSO and OptoIso. Average curves were plotted using cells from 4 independent experiments. **(B)** HeLa cells expressing β1AR-CFP, cAMPr, and Nanobody80-mCherry were treated with 100 μM OptoIso and incubated for 30 minutes. Then the cell culture dish was washed with HBSS 5 times. Following that the cells were exposed to blue light while imaging. cAMPr fluorescence increase and the Nb80-mCherry recruitment only to the endomembranes was observed. The line plot and the whisker box plot show the cytosolic cAMPr dynamics compared to the basal level with OptoIso and DMSO, upon blue light exposure. Average curves were plotted using cells from 3 independent experiments. The error bars represent SD (standard deviation of mean). The scale bar = 5 µm. CFP: Cyan Fluorescent Protein; OptoIso: Tri protected Isoproterenol; Cyto: cytosolic; DMSO: Dimethyl Sulfoxide; Nb80: Nanobody80; mCh: mCherry. The blue box indicates the blue light exposure. The yellow-colored arrows indicate the endomembrane β1AR expression. Green box indicates the Venus-Gγ9 photobleaching. Blue box indicates the blue light exposure.

We next examined whether such Gβγ mobility increases could be used to measure endomembrane exclusive heterotrimer activation. Similar to the above cell surface β1AR experiment, cells were prepared and treated with 100 µM of OptoIso for 30 minutes before washing them with HBSS 5 times. To determine pre-β1AR activation Gγ9 shuttling, we photobleached Venus-Gγ9 in the endomembrane regions indicated by the β1AR-CFP expression and measured FRAP t_1/2_ (Fig. 8A, green box). To determine Gγ9 shuttling after β1AR activation, we exposed cells to blue light until the Nb80 recruitment to the endomembrane reached equilibrium. Next, we similarly and selectively photobleached Venus on the endomembranes and measured FRAP t_1/2_. We observed a 2-fold faster FRAP t_1/2_ after OptoIso-induced β1AR activation (Fig. 8A-whisker box plot). One-way ANOVA showed that the mobilities of Gγ9 before the optical activation of OptoIso are significantly different from that of the after optical activation (Fig. 8A-Whisker box plots, one-way ANOVA: *F_1,21_* = 20.77, *p* = 1.7143, Table S2A and B). Control cells treated with DMSO (in place of OptoIso) didn’t show a significant difference in before and after β1AR activation Gγ9 mobilities (One-way ANOVA: *F_1,16_* = 0.0196, *p* = 0.890, Table S3A and B).

We also examined whether blue light deprotection of OptoIso and subsequent β1AR activation at the endomembrane could stimulate cAMP generation. HeLa cells expressing β1AR-CFP, cAMPr (cAMP sensor), and Nb80-mCherry were first treated with 100 μM of OptoIso for 30 minutes and washed with HBSS 5 times. We next imaged the cells using 488 nm excitation/515 nm emission to capture cAMPr fluorescence at 1-second intervals. To activate OptoIso, we initiated blue light exposure at t=50 seconds. The cells showed significant cAMPr fluorescence increases upon blue light (Fig. 8B). The underlying β1AR activation was indicated by Nb80-mCherry recruitment to the endomembranes. The control cells, which were treated with DMSO, showed no cAMPr fluorescence increase or Nb80-mCherry recruitment upon blue light exposure (Fig. S5D). These data collectively indicate that OptoIso is a highly useful photo-pharmacology tool to optically activate endomembrane exclusive β1ARs and probe the resultant signaling at the proximity to the deeper cell organelles, including the nucleus.

Given the difficulty of exclusively controlling endomembrane GPCRs without the signaling contamination from the cell surface counterparts, we believe OptoIso provides several unprecedented advantages. Most importantly, it provides an absolute certainty of the prevention of signaling contamination from the plasma membrane receptors. The OptoIso-induced endomembrane receptor activation is not limited by the kinetics of cell-permeable ligand diffusion but only by photoactivation kinetics. Additionally, OptoIso allows us to activate sub-endomembrane regions of single cells with spatiotemporal control, which in turn gives us the ability to make a fair comparison of single-cell behaviors in a cell population. Further, OptoIso is not deprotected by green, yellow, and red wavelengths, and therefore, sensors with multiple fluorescence tags can be used to map the dynamics of several signaling molecules simultaneously while optically activating βAR signaling.

## 3. MATERIALS AND METHODS

### 3.1 Reagents

The reagents used were as follows: Isoproterenol hydrochloride (Tocris Bioscience), dobutamine (Cayman chemical), sotalol (Cayman chemical), and DMSO (Focus Biomolecules). Reagents and solvents used in the synthesis of OptoIso, DiIsoprot, and MonoIsoprot were acquired from Oakwood Chemicals (reagents) or Sigma-Aldrich (solvents). According to the manufacturer’s instructions, all the reagents were dissolved in appropriate solvents and diluted in 1% Hank’s balanced salt solution supplemented with NaHCO_3_, or a regular cell culture medium, before adding to cells.

### 3.2 preparation of Triprotected Isoproterenol

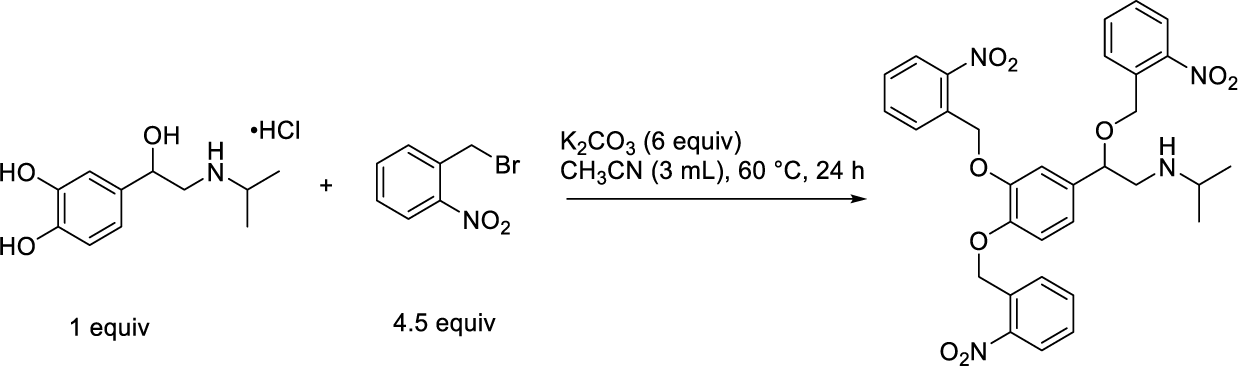

A 7.5 mL vial was charged with Isoproterenol (100 mg, 0.40 mmol, 1.0 equiv.), 2-nitrobenzyl bromide (400 mg, 1.82 mmol, 4.5 equiv), K_2_CO_3_ (331 mg, 2.4 mmol, 6.0 equiv.), and acetonitrile (3 mL). The vial was sealed with a PTFE lined cap and the reaction mixture was heated in a pie-block at 60 °C with stirring for 24 h. After cooling to room temperature, the solvent evaporated *in vacuo*. The reaction mixture was then purified using Flash chromatography (25 % EtOAc in Hexane), and the product recovered as a yellow solid (177 mg, 72% yield). TLC (Hex: EtOAc= 7:3, R_f_ = 0.35).

### 3.3 Preparation of Diprotected Isoproterenol

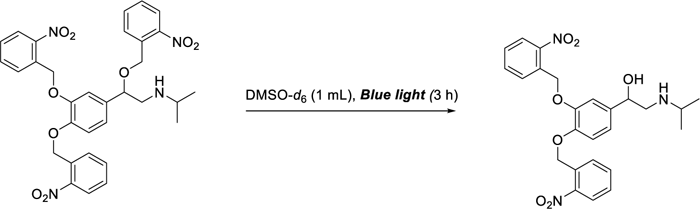

An NMR tube was charged with Triprotected Isoproterenol (90 mg, 0.146 mmol, 1 equiv.), followed by dissolving in DMSO-*d*_6_ (1 mL). The reaction was irradiated with a Kessil A160WE Tuna Blue lamp (in blue mode) for 3 h, until ^1^H NMR showed complete deprotection of the benzylic ether. The reaction mixture was then diluted with water and extracted with EtOAc (3 x 10 mL). The product was further purified using a pipette column (eluted with DCM, EtOAc, and finally with MeOH). Product recovered as yellow oil (40 mg, 57% yield), which was dissolved in deuterated MeOH for NMR. TLC (MeOH: EtOAc= 20:80, R_f_ = 0.15).

### 3.4 Preparation of Monoprotected Isoproterenol

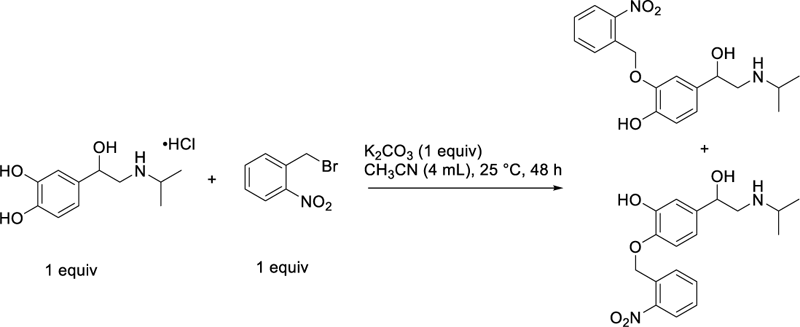

A 7.5 mL vial was charged with Isoproterenol (100 mg, 0.40 mmol, 1.0 equiv.), 2-nitrobenzyl bromide (82 mg, 0.40 mmol, 1.0 equiv), K_2_CO_3_ (1.0 equiv.), and acetonitrile (4 mL). The vial was sealed with a PTFE lined cap and the reaction mixture was stirred at 25 °C for 48 h. Afterwards the solvent was evaporated *in vacuo*, and the residue was filtered through a silica plug, first eluting with EtOAc, then with EtOAc: MeOH. The partially purified mixture which contained both di and monoprotected Isoproterenol was then purified using automated Flash chromatography (Combi-Flash by Teledyne) over C18 silica (Gradient 0% MeOH/Water -> 50% MeOH/Water), and product recovered as a light brown oil containing a mixture of monoprotected catecholamines (15.6 mg, 11% yield).

### 3.5 DNA constructs

DNA constructs used were as follows: β1AR-CFP, Venus-miniGs, Venus-Gγ9, Gαs-YFP, cAMPr, Nanobody80-mCherry. Venus-miniGs is a kind gift from Dr. Nevin Lambert, Augusta University, GA. Nanobody 80 (Nb80) was kindly provided by Dr. Roshanak Irannejad (University of California, San Francisco, CA). β1AR-CFP, Venus-Gγ9, and Gαs-YFP are a gift from Dr. N. Gautam’s laboratory, Washington University in St Louis, MO. All cloning was performed using Gibson assembly cloning (NEB). All cDNA constructs were confirmed by sequencing.

### 3.6 Cell culture and DNA transfection

HeLa cells were purchased from ATCC, USA. Recommended cell culture media; HeLa (MEM/10%DFBS/1%PS) were used to subculture cells on 29 mm, 60 mm, or 100 mm cell culture dishes. For live-cell imaging experiments, cells were seeded on 29 mm glass-bottomed dishes at a density of 1 x 10^5^ cells.

DNA transfections were performed using Lipofectamine^®^ 2000 reagent according to the manufacturer’s recommended protocols.

### 3.7 Live cell imaging, image analysis, and data processing

The methods, protocols, and parameters for live cell imaging are adapted from previously published work.^38, 64, 65^ Briefly, live cell imaging experiments were performed using a spinning disk confocal imaging system (Andor Technology) with a 60X, 1.4 NA oil objective, and iXon ULTRA 897BVback-illuminated deep-cooled EMCCD camera. Photoactivation and Spatio-temporal light exposure on cells in regions of interest (ROI) was performed using a laser combiner with a 445 nm solid-state laser delivered using Andor® FRAP-PA (fluorescence recovery after photobleaching and photoactivation) unit in real-time, controlled by Andor iQ 3.1 software (Andor Technologies, Belfast, United Kingdom). Fluorescent proteins such as Nanobody 80-mCherry were imaged using 594 nm excitation−624 nm emission settings. Venus-miniGs, Venus-Gγ9 and Gαs-YFP were imaged using 515 nm excitation and 542 nm emission. β1AR-CFP was imaged using 445 nm excitation and 478 nm emission. For global and confined optical activation of OptoIso, 445 nm solid-state laser coupled to FRAP-PA was adjusted to deliver 145 nW power at the plane of cells, which scanned light illumination across the region of interest (ROI) at 1 ms/μm^2^. The time-lapse images were analyzed using Andor iQ 3.2 software by acquiring the mean pixel fluorescence intensity changes of the entire cell or selected area/regions of interest (ROIs). Briefly, the background intensity of images was subtracted from the intensities of the ROIs assigned to the desired areas of cells (plasma membrane, internal membranes, and cytosol) before intensity data collection from the time-lapse images. The intensity data from multiple cells were opened in Excel (Microsoft office®) and normalized to the baseline by dividing the whole data set by the average initial stable baseline value. Data were processed further using Origin-pro data analysis software (OriginLab®).

### 3.8 Designing of OptoIso model

In this study, we employed an integrative computational approach to design OptoIso using already available protein-ligand interaction structural information on β1-AR, combining molecular docking and advanced modeling techniques.^66^ Briefly, Isoproterenol bound β1-AR (PDB ID: 7JJO)^39^ structure was incorporated into the Schrodinger software, then we manually remove crystallization artifacts and irrelevant amino acid chains. Then the structure was optimized using the protein preparation tool in the Schrodinger Maestro software. We used this optimized structure to map the significant interactions between the ligand and the ligand binding pocket of the β1-AR. Then based on the interactions we designed three ligands masking the hydroxyl groups with photolabile groups using ChemDraw. After that we created glide docking grids on optimized β1-AR using bound Isoproterenol as the center using glide grid generation option in Schrodinger Maestro.^67, 68^ We used LigPrep tool in Schrodinger Maestro to prepare the ChemDraw structures of the Isoproterenol analogs and to model them with precise molecular characteristics. Then they were docked into the glide grid generated to examine their binding to the receptor in silico.^69^

### 3.9 Experimental rigor and Statistical analysis

To eliminate potential biases or preconceived notions and improve the experimental rigor, we used the reagent-blinded-experimenter approach for the key findings of our study. All experiments were repeated multiple times to test the reproducibility of the results. Results are analyzed from multiple cells and represented as mean±SD. The exact number of cells used in the analysis is given in respective figure legends. Digital image analysis was performed using Andor iQ 3.1 software, and fluorescence intensity obtained from regions of interest was normalized to initial values (baseline). Data plot generation and statistical analysis were done using OriginPro software (OriginLab®). One-way ANOVA statistical tests were performed using OriginPro to determine the statistical significance between two or more populations of signaling responses. Tukey’s mean comparison test was performed at the p < 0.05 significance level for the one-way ANOVA statistical test. No unexpected or unusually high safety hazards were encountered throughout the experimental procedures in this manuscript.

### 3.10 Nuclear magnetic resonance spectroscopy experiments

Analysis was performed on a Varian Inova 600 MHz Spectrometer. Data was processed using MestReNova software.*3.11 High resolution mass spectrometric experiments*.

### 3.11 High resolution mass spectrometric experiments

Analysis was performed on a Q-Exactive Orbitrap high resolution mass spectrometer (HRMS) from Thermo Fisher Scientific (Waltham, MA) for detection and fragmentation experiments. A Thermo Scientific autosampler injector was used for the direct injection for experiments with the Vanquish UHPLC 202 system. A 50 μm internal diameter (i.d.) and 365 μm outer diameter (o.d) capillary from Polymicro Technologies (Phoenix, AZ) was used as connection tubing between the auto-sampler and the Thermo nanospray Flex ion source emitter in positive ionization mode according to the manufacturer guidelines. To improve ionization,^70^ a nano emitter with a split flow system was created.^71^

Briefly, a window was generated in a 20 cm long fused silica 50 μm i.d., 365 o.d. capillary using an electrical arc to remove 0.5 cm of the polyimide coating. Photopolymerized frits were generated using a monomer mix of 350 μL trimethylolpropane trimethacrylate and 150 μL of glycidyl methacrylate with 7.9 mg of benzoin methyl ether (BME). The porogenic solvent was prepared by mixing 250 μL of toluene and 750 μL of isooctane. The monomer solution (300 mL) was added to the porogen solution and sonicated for 15 min. The frit mixture is loaded into the capillary and polymerization was initiated with UV-lamp (UVP, Cambridge, U.K.); wavelength was 365 nm, 6 W, 0.12 A, time for the reaction was 30 min at ambient temperature. These frit keeps a stable pressure and flow and prevent debris from entering to the mass spectrometer.

In order to create a nano emitter, tips were generated using a laser puller model P-2000 (Sutter Instruments, Novato, CA) with heating time 420 ms, velocity 80 ms, delay time 150 ms, and pulling time 225 ms. The nano emitter fritted capillary was etched in hydrofluoric acid (51%) to open a fine tip resulting in the nanospray emitter. The nanospray voltage was at 2 kV spray voltage, 150 °C mass spectrometry ion transfer tube capillary temperature. A T-splitter was used to carry the inlet flow from the auto sampler and outlet was split to a 50 cm open tube capillary of 75 μm i.d., 365 o.d. to split to waste and the other end was for the created nano emitter electro spraying the effluent to the HRMS. Direct injection of 1 μL sample volume in a mixed scan modes, full scan, and fragmentation scans. Fragmentation using steeped collision energy was done on 25, 35, 45, and 55 % on 0.4 u isolation window. Spectra were collected at a mass range of 70-1000 m/z with a mass resolution of 70,000 (RFWHM) and automatic gain control (AGC) of 1E6. OptoIso synthesized dry compound was dissolved in methanol for HRMS injections (1 μL injection volume). The data analysis and fragment generation were determined and presented using ChemDraw.

## Supporting information

Thotamune et al- Supporting Information

## Acknowledgment

We acknowledge Dr. N. Gautam (Washington University-School of Medicine, St. Louis, MO, USA) and Dr. Nevin Lambert (Augusta University, GA) for providing us with plasmid DNA. We also thank Dr. Roshanak Irannejad (University of California, San Francisco, CA). for providing us with plasmid DNA. We acknowledge Dr. James L. Edwards for kindly providing us the HRMS facilities and reviewing the manuscript. We also thank Ajith lab members for various experimental support and discussions. We thank the Saint Louis University Institute for Drug and Biotherapeutic Innovation for providing computational resources and access to Schrödinger software with funding from the Saint Louis University Research Institute. We thank Dr. Yongwah Kim (The University of Toledo, OH) for helpful discussions related to our NMR experiments.

## Abbreviations

GPCRs: (G protein coupled receptors)
β1AR: (β adrenergic receptor)
GTP: (Guanosine-5’-triphosphate)
PKC: (protein kinase C)
Nt: (N terminus)
Ct: (C terminus)
GRK: (G protein-coupled receptor kinase)
HRMS: (high resolution mass spectrometer)

## Author contribution

W.T. and S.U. conducted the majority of experiments and performed the data analysis. K.K.S. and M.C.Y. synthesized the protected isoproterenol derivatives, performed NMR and studied Photodeprotection under photolysis. W.T performed computational docking and bioinformatics analysis. M.E.M. performed HRMS analysis. W.T. and S.U. performed the statistical analysis. A.K., M.C.Y., W.T., and S.U. conceptualized the project. A.K, M.C.Y, W.T, S.U and K.K.S wrote and reviewed the manuscript.

## Data availability statement

The datasets used and analyzed during the current study are available from the corresponding author upon reasonable request.

