## Supplementary material for "Optical Control of Cell-Surface and Endomembrane-Exclusive β-Adrenergic Receptor Signaling": Thotamune et al- Supporting Information

**Figure S1**

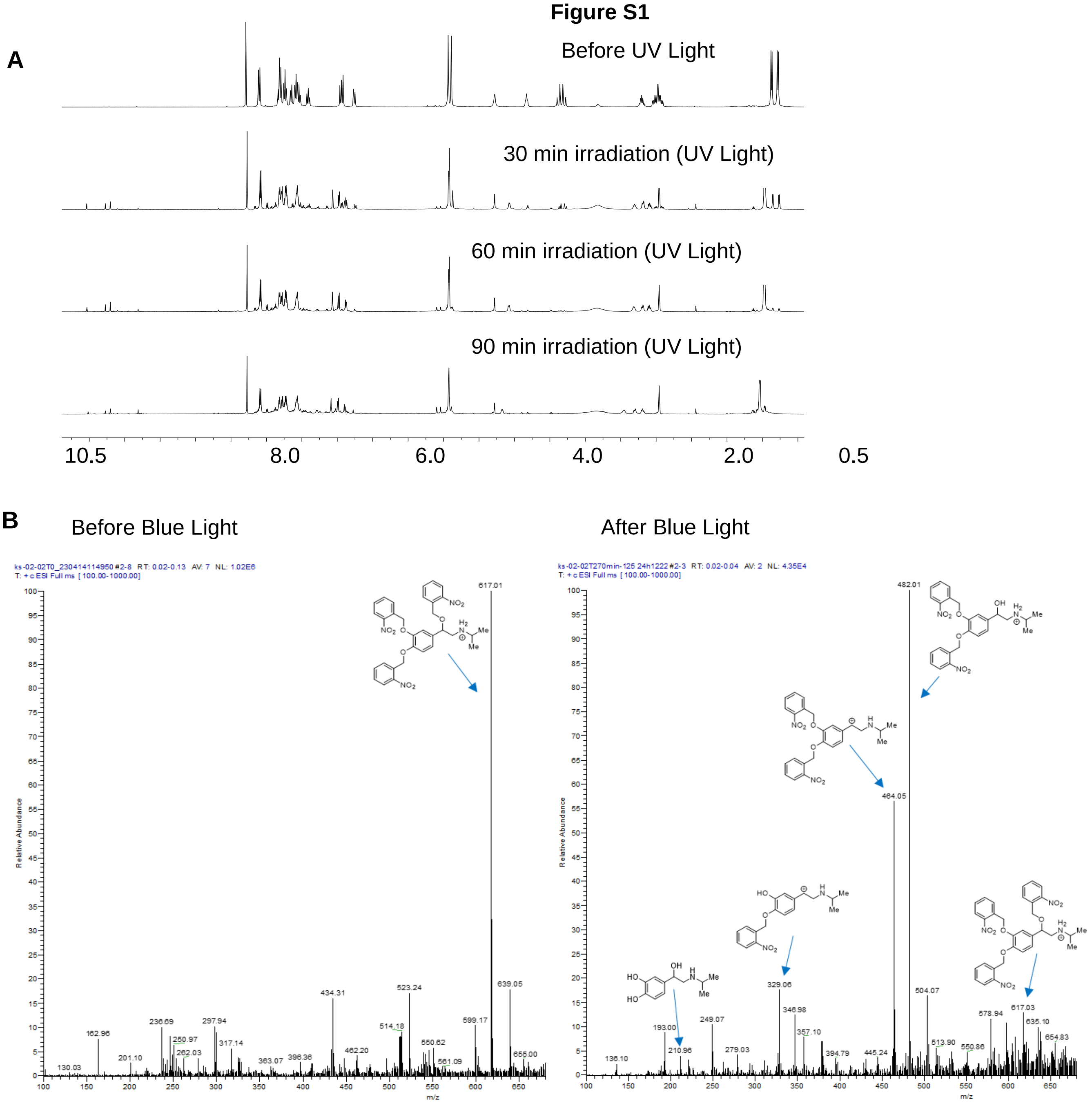

**Figure S1. (A)** $^{1}H$ NMR Analysis of the Photodeprotection of TriIsoprot using a Spectroline E-Series (DMSO-$d_6$, 600 MHz, 298 K). **(B)** ESI-MS spectrums of OptoIso before and after 15 minutes of blue light exposure. Photo-deprotection of the OptoIso generated enough potent ligands to activate the receptors.

### Figure S2

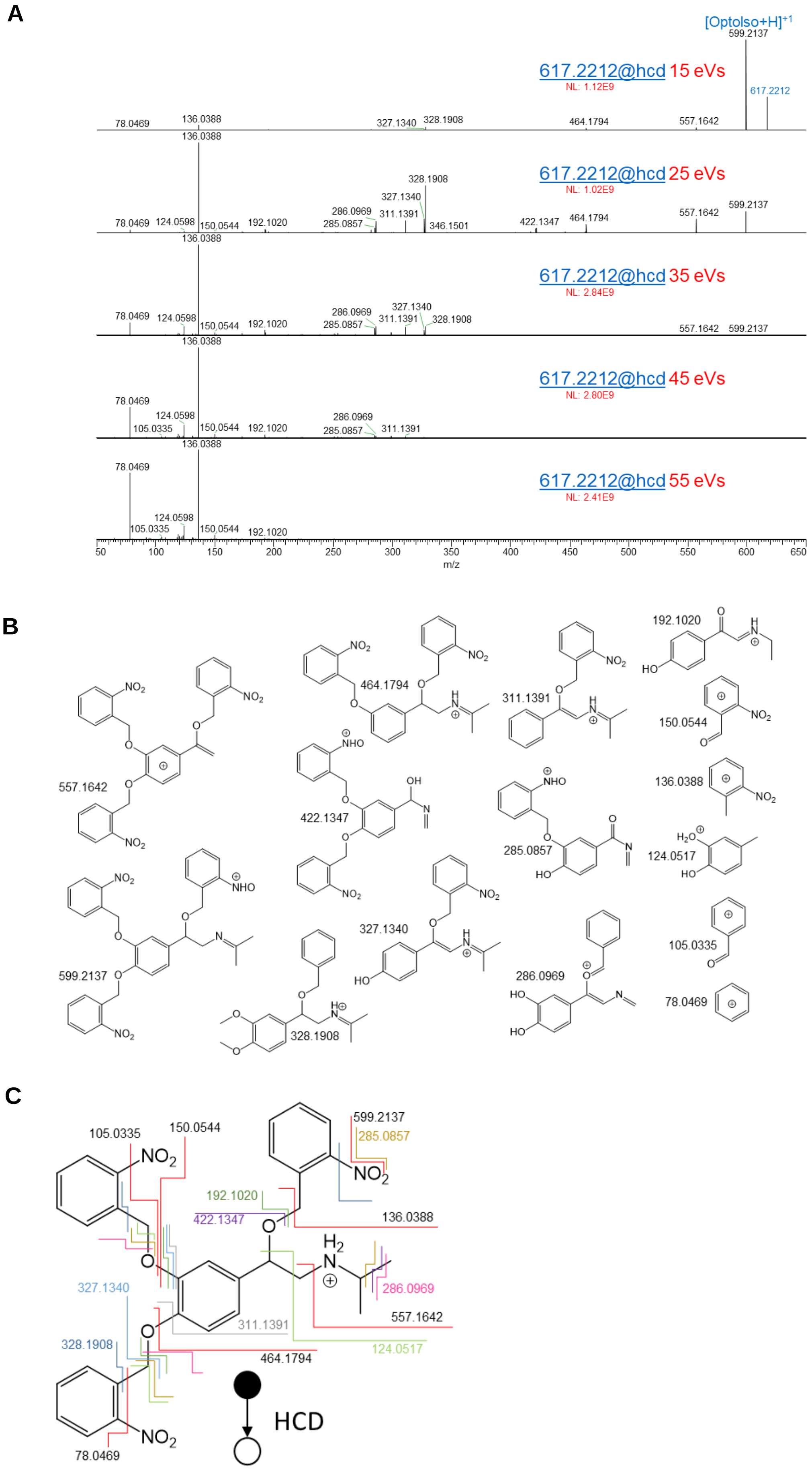

A
[Optolso+H]+1
617.2212@hcd 15 eVs
NL: 1.12E9
617.2212@hcd 25 eVs
NL: 1.02E9
617.2212@hcd 35 eVs
NL: 2.84E9
617.2212@hcd 45 eVs
NL: 2.80E9
617.2212@hcd 55 eVs
NL: 2.41E9
78.0469
105.0335
124.0598
136.0388
150.0544
192.1020
285.0857
286.0969
311.1391
327.1340
328.1908
346.1501
422.1347
464.1794
557.1642
599.2137
617.2212
m/z
B
557.1642
464.1794
311.1391
192.1020
150.0544
136.0388
422.1347
285.0857
124.0517
599.2137
327.1340
328.1908
286.0969
105.0335
78.0469
C
105.0335
150.0544
599.2137
285.0857
192.1020
422.1347
136.0388
327.1340
286.0969
311.1391
557.1642
328.1908
124.0517
464.1794
78.0469
HCD

**Figure S2. (A)** The fragmentation HRMS spectrums ($MS^2$) of the OptoIso upon subjecting the sample steeped collision energy of 15, 25, 35, 45, and 55 electron volts (eVs). The $MS^2$ fragments match the expected cleavage points in Figure S1C. OptoIso and all fragments are protonated in the positive ionization mode as analyte (M) with a proton (H) with a charge state of 1 (i.e. $[M+H]^{+1}$). NL is the normalized level of signal found in the HRMS fragmentation spectrums. **(B)** OptoIso fragments upon fragmentation and their corresponding exact mass. **(C)** The chemical structure of the synthesized purified tri protected Isoproterenol (OptoIso). The structure was drawn by ChemDraw® Perkin Elmer™. HCD is the high-energy collision dissociation by colliding orbit-rapped ions with nitrogen gas. The colored lines represent the fragmentation cleavage points. Multiple lines of the same color correspond to multiple fragmentation events to produce the fragment. The numbers are the exact mass of the found fragments upon collision with nitrogen gas inside the HRMS-Orbitrap.

**Figure S3**

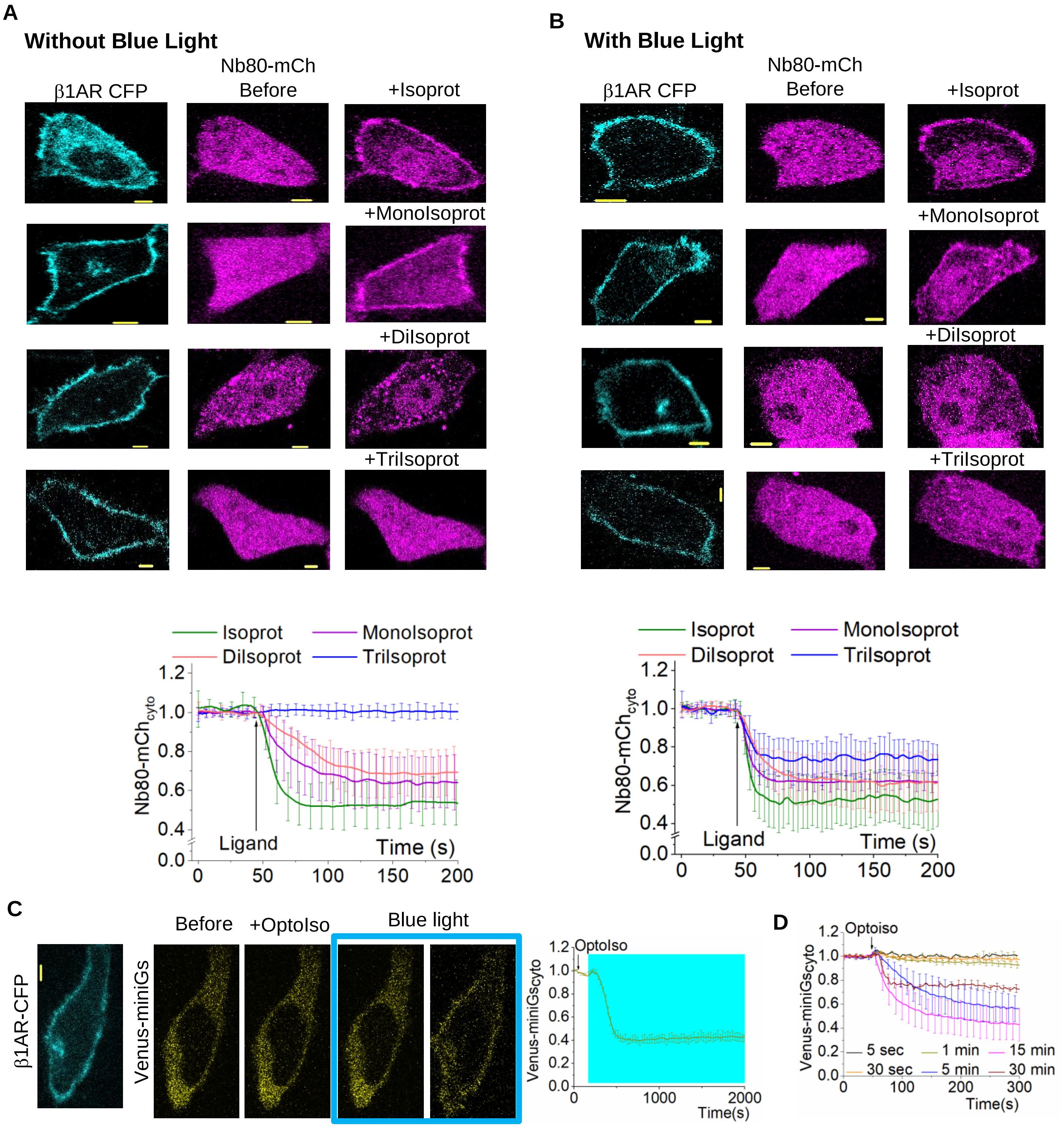
A
Without Blue Light
β1AR CFP
Nb80-mCh
Before
+Isoprot
+MonoIsoprot
+DiIsoprot
+TriIsoprot
B
With Blue Light
β1AR CFP
Nb80-mCh
Before
+Isoprot
+MonoIsoprot
+DiIsoprot
+TriIsoprot
Isoprot
MonoIsoprot
DiIsoprot
TriIsoprot
Nb80-mCh cyto
1.2
1.0
0.8
0.6
0.4
0.0
Ligand
Time (s)
0
50
100
150
200
C
β1AR-CFP
Venus-miniGs
Before
+OptoIso
Blue light
1.2
1.0
0.8
0.6
0.4
0.2
0.0
OptoIso
Venus-miniGscyto
0
1000
2000
Time(s)
D
Optoiso
Venus-miniGscyto
5 sec
30 sec
1 min
5 min
15 min
30 min
0
100
200
300
Time(s)

**Figure S3. (A)** HeLa cells expressing β1AR-CFP and Nanobody80-mCherry were treated either with 100 μM of Isoproterenol or three protected isoproterenol variants; Mono protected (MonoIsoprot), diprotected (DiIsoprot) and tri-protected (TriIsoprot) at 50 seconds (all the compounds are not exposed to blue light). Isoproterenol treated cells showed robust Nanobody80-mCherry recruitment to the plasma membrane and MonoIsoprot and DiIsoprot-treated cells showed a detectable Nanobody80 translocation to the plasma membrane. OptoIso-treated cells didn't show detectable Nanobody80-mCherry translocation to the plasma membranes of the cells. The plot shows the cytosolic Nanobody80-mCherry dynamics normalized to the basal level fluorescence. **(B)** HeLa cells expressing β1AR-CFP and Nanobody80-mCherry were treated either with 100 μM of Isoproterenol or three protected isoproterenol variants: Mono protected (MonoIsoprot), deprotected (DiIsoprot) and tri-protected (OptoIso) at 50 seconds, after exposing the compounds to blue light for 15 minutes. Isoproterenol-treated cells showed robust Nanobody80-mCherry recruitment to the plasma membrane and MonoIsoprot and DiIsoprot-treated cells showed a detectable Nanobody80-mCherry translocation to the plasma membrane. OptoIso-treated cells also showed significant Nanobody80-mCherry translocation to the plasma membranes of the cells. The plot shows the cytosolic Nanobody80-mCherry dynamics normalized to the basal level fluorescence. **(C)** HeLa cells expressing β1AR-CFP, Venus-miniGs were exposed to 100 μM OptoIso at 50 seconds and blue light exposure was started at 3 minutes. Blue light exposure induced a gradual miniGs recruitment to the plasma membrane. **(D)** HeLa cells expressing β1AR-CFP, Venus-miniGs were exposed to OptoIso aliquots with different blue light exposure times. Average curves were plotted using cells from ≥3 independent experiments. The error bars represent SD (standard deviation of mean). The scale bar = 5 μm. CFP: Cyan Fluorescent Protein; MonoIsoprot: Monoprotected Isoproterenol; DiIsoprot: Di protected Isoproterenol; OptoIso: Tri protected Isoproterenol; Cyto: cytosolic.

**Figure S4**

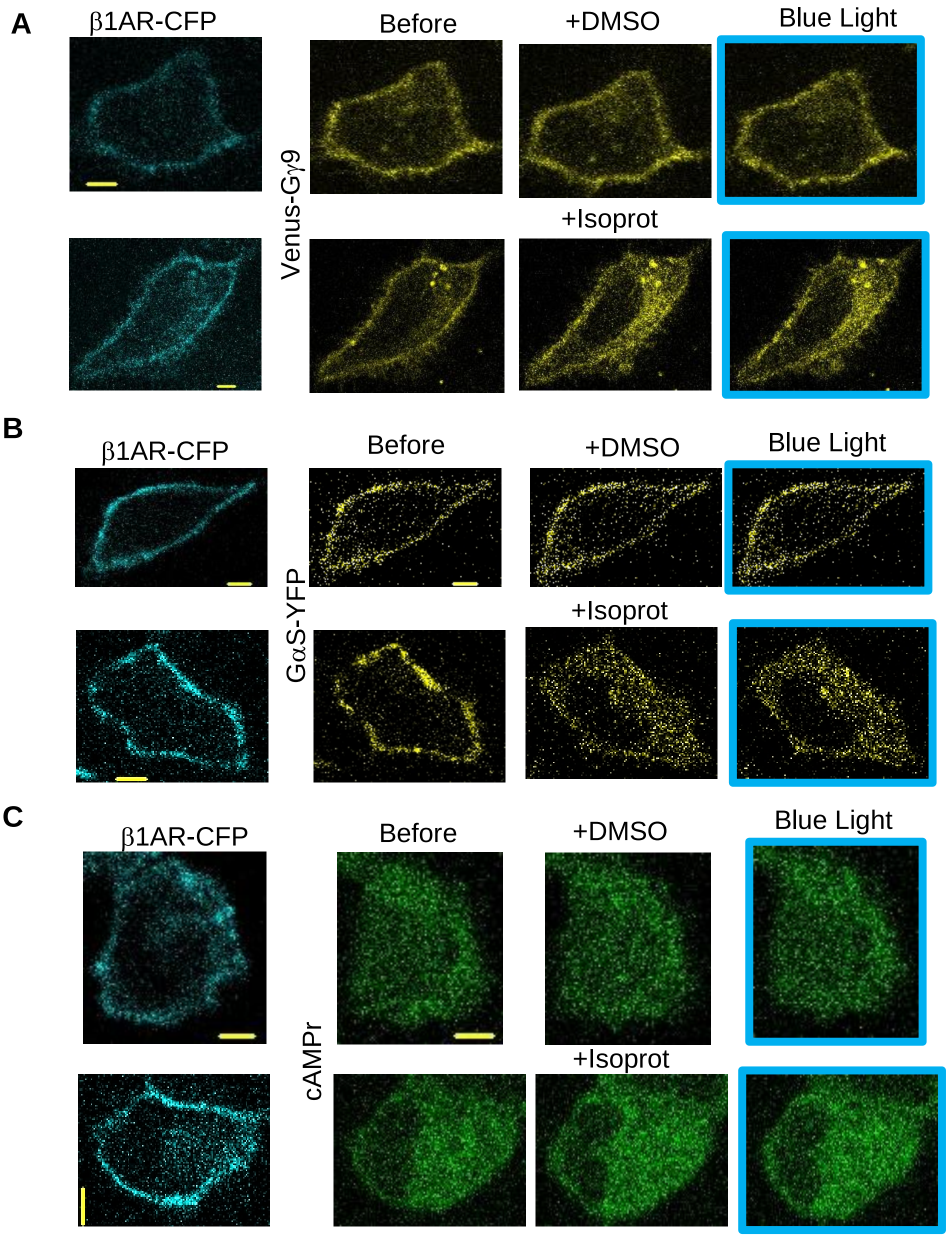

A
β1AR-CFP
Before
+DMSO
Blue Light
Venus-Gγ9
+Isoprot
B
β1AR-CFP
Before
+DMSO
Blue Light
GαS-YFP
+Isoprot
C
β1AR-CFP
Before
+DMSO
Blue Light
cAMPr
+Isoprot

**Figure S4. (A)** HeLa cells expressing β1AR-CFP and Venus-Gγ9 were treated with DMSO at 50 seconds and exposed to blue light at 3 minutes. The cells didn't show a detectable Gβγ translocation to the endomembranes. Then the same cells were treated with 100 μM of Isoproterenol at 50 seconds and blue light was given at 3 minutes. The cells showed robust Gβγ translocation immediately after ligand addition. **(B)** HeLa cells expressing β1AR-CFP and Gαs-YFP were treated with DMSO at 50 seconds and exposed to blue light at 3 minutes. The cells didn't show a detectable Gαs-YFP translocation to the cytosol. Then the same cells were treated with 100 μM of Isoproterenol at 50 seconds and blue light was given at 3 minutes. The cells showed robust Gαs translocation upon ligand addition. **(C)** HeLa cells expressing β1AR-CFP and cAMPr (cAMP sensor) were treated with DMSO at 50 seconds and exposed to blue light at 3 minutes. The cells didn't show a detectable cAMPr fluorescence increase. Then the same cells were treated with 100 μM of Isoproterenol at 50 seconds and blue light was given at 3 minutes. The cells showed robust cAMPr fluorescence increase upon ligand addition. The error bars represent SD (standard deviation of mean). The scale bar = 5 μm. CFP: Cyan Fluorescent Protein; IP: isoproterenol; DMSO: Dimethyl Sulfoxide. The blue box indicates the blue light exposure.

**Figure S5**

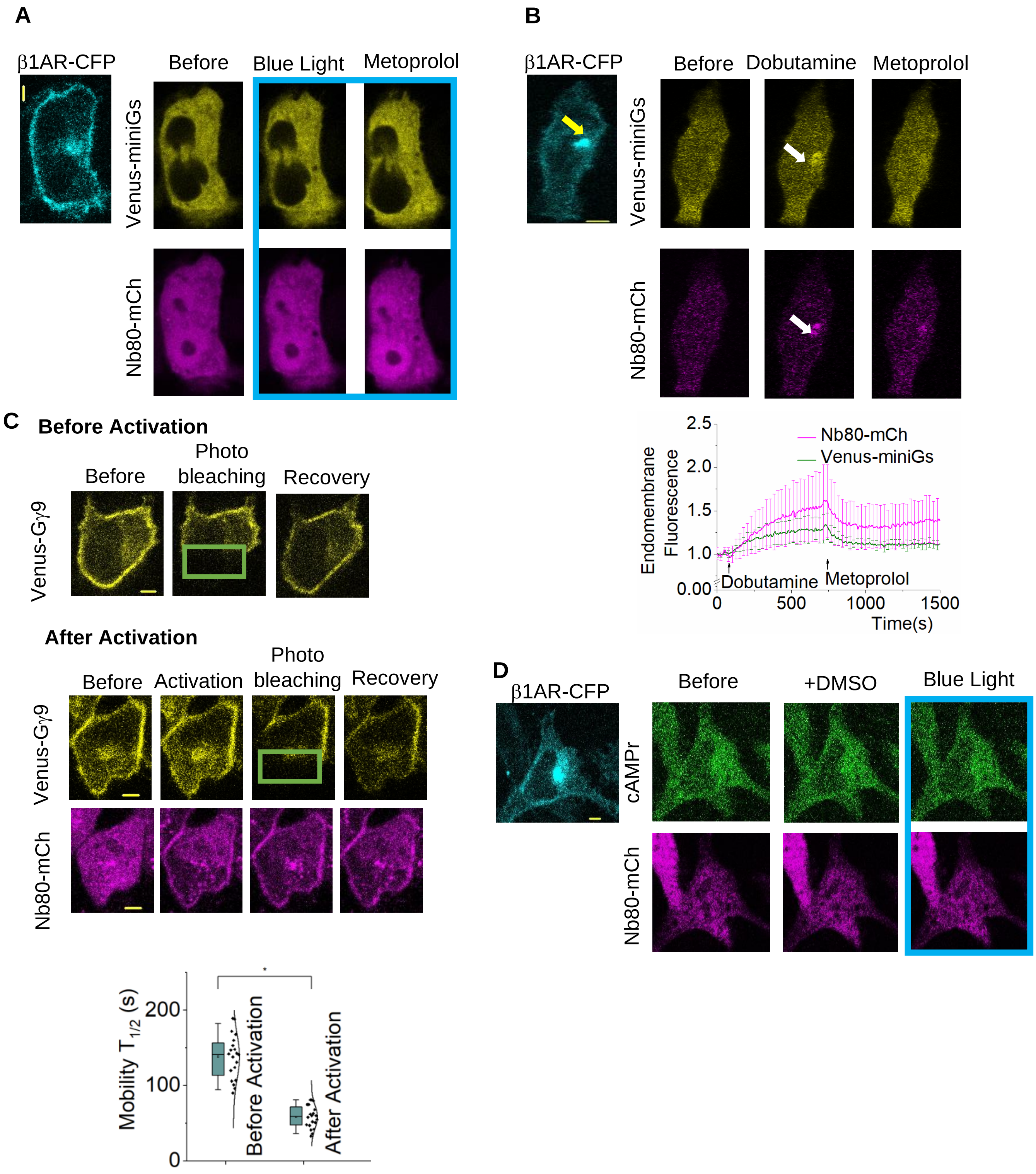
A
β1AR-CFP
Before
Blue Light
Metoprolol
Venus-miniGs
Nb80-mCh
B
β1AR-CFP
Before
Dobutamine
Metoprolol
Venus-miniGs
Nb80-mCh
Nb80-mCh
Venus-miniGs
Endomembrane Fluorescence
2.5
2.0
1.5
1.0
0.00
Dobutamine
Metoprolol
0
500
1000
1500
Time(s)
C
Before Activation
Before
Photo bleaching
Recovery
Venus-Gγ9
After Activation
Before
Activation
Photo bleaching
Recovery
Venus-Gγ9
Nb80-mCh
Mobility $T_{1/2}$ (s)
200
100
0
Before Activation
After Activation
D
β1AR-CFP
Before
+DMSO
Blue Light
cAMPr
Nb80-mCh

**Figure S5. (A)** HeLa cells expressing β1AR-CFP, Venus-miniGs and Nanobody80-mCherry were treated with 10 μM dobutamine A robust Venus-miniGs and Nanobody80-mCherry recruitment were observed exclusively to the endomembranes inside the cells. The disappearance of these sensors from the endomembranes was observed at 12 minutes, upon addition of 10 μM Metoprolol. The plots show the Venus- MiniGs and Nanobody80-mCherry dynamics at endomembranes. **(B)** HeLa cells expressing β1AR-CFP, Venus-miniGs and Nanobody80-mCherry were treated with DMSO and incubated for 30 minutes. Then the cell culture dish was washed with HBSS for 5 times. The cells didn't exhibit detectable Venus-miniGs or nanobody80-mCherry recruitment to the endomembranes. **(C)** β1AR-CFP, Venus-Gγ9 and Nanobody80-mCherry were expressed in HeLa cells. Venus- Gγ9 on the plasma membranes were photobleached and the fluorescence recovery after photobleaching was examined before the receptor activation. Nanobody80-mCherry was cytosolic since the receptors were not activated. Then the cells were exposed 100 μM Isoprot and Nanobody80-mCherry recruitment was observed at the plasma membranes. Then Venus- Gγ9 was photobleached and the fluorescence recovery after photobleaching was examined. The whisker box plots show mobility half-time ($t_{1/2}$) of Venus- Gγ9 before and after the receptor activation. **(D)** HeLa cells expressing β1AR-CFP, cAMPr and Nanobody80-mCherry were treated with DMSO and incubated for 30 minutes. Then the cell culture dish was washed with HBSS for 5 times. The cells didn't show nanobody80-mCherry recruitment or cAMPr fluorescence increase upon blue light exposure. The scale bar = 5 μm. CFP: Cyan Fluorescent Protein; DMSO: Dimethyl Sulfoxide; Nb80: Nanobody80; mCh: mCherry. The blue box indicates the blue light exposure.

Table S1-A: One-way ANOVA statistics for mobility $T_{1/2}$ of Gγ9 before and after activation of PM bound β1ARs with Isoproterenol.

| Descriptive Statistics | | | | |
| --- | --- | --- | --- | --- |
| | N Analysis | Mean | Standard Deviation | SE of Mean |
| Before activation | 20 | 152.2 | 48.65301 | 10.87914 |
| After activation | 20 | 58.4 | 14.92931 | 3.33829 |

Table S1-B:

| Overall ANOVA | | | | | |
| --- | --- | --- | --- | --- | --- |
| | DF | Sum of Squares | Mean Square | F value | Prob>F |
| Model | 1 | 87984.4 | 87984.4 | 67.94162 | <0.0001 |
| Error | 38 | 49210 | 1295 | | |
| Total | 39 | 137194.4 | | | |

At the 0.05 level, the population means are significantly different.

Table S2-A: One-way ANOVA statistics for mobility $T_{1/2}$ of G$\gamma$9 before and after activation of endomembrane exclusive $\beta$1ARs using OptoIso.

| Descriptive Statistics | | | | |
|---|---|---|---|---|
| | N Analysis | Mean | Standard Deviation | SE of Mean |
| Before activation | 11 | 33.84439 | 9.72369 | 2.9318 |
| After activation | 12 | 16.39671 | 8.63891 | 2.49384 |

Table S2-B:

| Overall ANOVA | | | | | |
|---|---|---|---|---|---|
| | DF | Sum of Squares | Mean Square | F value | Prob>F |
| Model | 1 | 1747.1143 | 1747.1143 | 20.77023 | 1.71436E-4 |
| Error | 21 | 1766.44133 | 84.11625 | | |
| Total | 22 | 3513.55564 | | | |

At the 0.05 level, the population means are significantly different.

Table S3-A: One-way ANOVA statistics for mobility $T_{1/2}$ of Gγ9 before and after blue light exposure for cells treated with DMSO.

| Descriptive Statistics | | | | |
|---|---|---|---|---|
| | N Analysis | Mean | Standard Deviation | SE of Mean |
| Before activation | 9 | 13.48556 | 3.15079 | 1.05026 |
| After activation | 9 | 13.30843 | 2.10727 | 0.70242 |

Table S3-B:

| Overall ANOVA | | | | | |
|---|---|---|---|---|---|
| | DF | Sum of Squares | Mean Square | F value | Prob>F |
| Model | 1 | 0.1412 | 0.1412 | 0.01965 | 0.89026 |
| Error | 16 | 114.94465 | 7.18404 | | |
| Total | 17 | 115.08585 | | | |

At the 0.05 level, the population means are not significantly different.

### SUPPLEMENTARY MOVIE LEGENDS

**Movie S1.** OptoIso induced reversible Venus-miniGs recruitment.

**Movie S2.** OptoIso induced reversible Nb80-mCherry recruitment.

**Movie S3.** Localized optical activation of OptoIso induced spatiotemporal Nb80-mCherry recruitment to the endomembranes (Blue box).

**Movie S4.** Localized optical activation of OptoIso induced spatiotemporal Venus-miniGs recruitment endomembranes (Blue box).
